# Dual-targeted nanoparticulate drug delivery systems for enhancing triple-negative breast cancer treatment

**DOI:** 10.1101/2024.01.23.576787

**Authors:** Shunzhe Zheng, Meng Li, Wenqian Xu, Jiaxin Zhang, Guanting Li, Hongying Xiao, Xinying Liu, Jianbin Shi, Fengli Xia, Chutong Tian, Ken-ichiro Kamei

## Abstract

The efficacy of DNA-damaging agents, such as the topoisomerase I inhibitor SN38, is often compromised by the robust DNA repair mechanisms in tumor cells, notably homologous recombination (HR) repair. Addressing this challenge, we introduce a novel nano-strategy utilizing binary tumor-killing mechanisms to enhance the therapeutic impact of DNA damage and mitochondrial dysfunction in cancer treatment. Our approach employs a synergistic drug pair comprising SN38 and the BET inhibitor JQ-1. We synthesized two prodrugs by conjugating linoleic acid (LA) to SN38 and JQ-1 via a cinnamaldehyde thioacetal (CT) bond, facilitating co-delivery. These prodrugs co-assemble into a nanostructure, referred to as SJNP, in an optimal synergistic ratio. SJNP was validated for its efficacy at both the cellular and tissue levels, where it primarily disrupts the transcription factor protein BRD4. This disruption leads to downregulation of BRCA1 and RAD51, impairing the HR process and exacerbating DNA damage. Additionally, SJNP releases cinnamaldehyde (CA) upon CT linkage cleavage, elevating intracellular ROS levels in a self-amplifying manner and inducing ROS-mediated mitochondrial dysfunction. Our results indicate that SJNP effectively targets murine triple-negative breast cancer (TNBC) with minimal adverse toxicity, showcasing its potential as a formidable opponent in the fight against cancer.

## 1. Introduction

Malignant tumors represent a formidable menace to human health, constituting one of the most arduous and ungratifying medical challenges [1, 2]. Amidst various approaches invented in the combat against cancer, genotoxic chemotherapeutic agents have ascended as principal interventions in the clinical management of diverse malignancies [3, 4]. A prime example of this is irinotecan, a topoisomerase I inhibitor, which has gained FDA approval for the treatment of colorectal cancer [5]. To elicit cytotoxic effects, irinotecan undergoes enzymatic conversion by carboxylesterases to yield its active form, 7-ethyl-10-hydroxy-camptothecin (SN38), which creates lethal double-strand breaks (DSBs) in DNA duplexes [6–7]. However, the effectiveness of such treatments is often undermined by the ability of tumor cells to repair these DSBs through HR, thereby regaining vitality and potentially developing drug resistance [8, 9].

Confronted with the challenges, a leading approach has emerged as modulating DNA damage responses (DDRs) to enhance the effectiveness of DNA-damaging agents [10]. Among diverse molecular targets, the bromodomain and extra-terminal (BET) protein family can intricately regulate the transcription of genes governing cell growth, cell cycle and differentiation [11, 12]. Especially, the BET number BRD4 regulates the expression of BRCA1 and Rad51 protein, which are recognized as pivotal components in various stages of HR, including lesion recognition, end resection, strand invasion and chain extension [13–16]. Consequently, BET inhibitors present an avenue for synergy with DNA toxic agents. However, the traditional approach of sequential or simultaneous administration of chemotherapeutic drugs often faces limitations due to differing pharmacokinetics and drug distributions, which can reduce the efficacy of combination therapies [17, 18].

The advent of nanoparticulate drug delivery systems (Nano-DDSs) has opened new avenues in tumor-targeting drug delivery, particularly for combination therapies [17, 19]. These systems can encapsulate antitumor agents inside the nanocarriers, extending their systemic circulation time and capitalizing on the enhanced permeability and retention (EPR) effect to passively target tumor tissues, thereby mitigating off-target toxicities [20–22]. Crucially, Nano-DDSs can ensure co-delivery of different drugs to tumor cells in optimal ratios, maximizing the synergistic effects [23, 24]. Tailored Nano-DDSs for the co-delivery of DNA-damaging agents and DNA repair inhibitors are thus a promising approach to overcome the limitations of current DNA damage therapies.

Furthermore, the tumor microenvironment (TME), characterized by high redox level due to excessive tumor cell metabolism, has inspired the development of multiple redox-responsive Nano-DDSs for tumor-specific drug release while minimizing nonspecific toxicities [25, 26]. Perturbing the redox balance in tumor cells, particularly by augmenting oxidative stress levels, has garnered prominence as a therapeutic option for promoting mitochondrial dysfunction and cell apoptosis [27, 28]. Hence, tailored redox-responsive Nano-DDSs, providing binary tumor-killing modality by manipulating DNA damage and repair pathways in parallel with perturbing the redox balance of the TME, will constitute a potent strategy in the combat against cancer.

Here we introduce a nano-strategy employing binary tumor-killing mechanisms for the treatment of triple-negative breast cancer (TNBC). To enable the rational co-delivery of SN38 and JQ-1, small-molecule prodrug assembled nanoparticles (PANP) is employed as the Nano-DDSs. This strategy involves the synthesis of SN38-LA and JQ-1-LA prodrugs, which co-assemble into SJNP nanoparticles. In the ROS-overexpressed TME, these nanoparticles release active SN38 and JQ-1, along with a by-product, cinnamaldehyde (CA), which further elevates cellular ROS levels. This synergistic action disrupts the HR process and induces ROS-mediated mitochondrial dysfunction, offering a robust approach to combat TNBC with minimal adverse toxicity.

## 2. Materials and methods

### 2.1 Materials

SN38, JQ-1 (carboxylic acid), and linoleic acid (LA) were procured from Bidepharm Co., Ltd. (Shanghai, China). DSPE-PEG_2k_ was sourced from AVT (Shanghai) Pharmaceutical Tech Co., Ltd. (Shanghai, China). Coumarin-6 and glutathione (GSH) were acquired from Shanghai Macklin Biochemical Technology Co., Ltd. (Shanghai, China). DiR, Hoechst 33342, DAPI and 3-(4,5-dimethylthiazol-2-yl)-2,5-diphenyltetrazolium bromide (MTT) were supplied by Meilun Biotechnology Co., Ltd. (Dalian, China). Hydrogen peroxide (H_2_O_2_) was provided by Shanghai Aladdin Biochemical Technology Co., Ltd. (Shanghai, China). The ROS assay kit, GSH content assay kit, JC-1 detection kit, and TUNEL assay kit were sourced from Beyotime Biotechnology Co., Ltd. (Shanghai, China). All other reagents employed in this study met at least analytical grade standards.

#### Antibodies

Phospho-Histone H2AX-S139 Rabbit mAb (Abclonal, AP0687), BRCA1 rabbit pAb (Abclonal, A11034), RAD51 rabbit pAb (Abclonal, A6268), Ki67 Rabbit pAb (Abclonal, A11390), Cy3 Goat Anti-Rabbit IgG (Abclonal, A5007) and FITC Goat Anti-Rabbit IgG (Abclonal, AS011).

#### Animals

The experimental animals were sourced from Liaoning Changsheng Biotechnology Co., Ltd (Benxi, China). Stringent adherence to the Guidelines for the Care and Use of Laboratory Animals characterized the meticulous execution of all animal experiments. The study secured essential approval from the Institutional Animal Ethical Care Committee (IAECC) of Shenyang Pharmaceutical University, under the sanctioned protocol number SYPU-IACUC-C2022-03-12-107.

### 2.2. Chemical synthesis

The details of chemical synthesis methodologies are delineated in the Supplementary Materials, including the synthesis protocols for the CT linkage, SN38-LA and JQ-1-LA.

### 2.3. Preparation of PANP and Dye-labeled PANP

PANP was prepared using the one-step nanoprecipitation technique as previously reported [23]. In a concise manner, an acetone solution of the prodrug, containing DSPE-PEG_2k_ (20%, w/w), was incrementally added to deionized water with robust stirring, while the acetone solvent was subsequently eliminated through vacuum rotary evaporation. For the formulation of co-assembled PANP, SN38-LA and JQ-1-LA were pre-mixed in an acetone solution at different molar ratios (10:1, 5:1 to 1:5 and 1:10, SN38-LA to JQ-1-LA). The dye-labeled PANP was prepared according the same procedure, with the addition of 5% dye (5%, coumarin 6 or DiR), and the excess dye was removed through dialysis in deionized water.

### 2.4 Characterization of PANP

The prepared PANP was diluted to a final concentration of 50 μg mL^−1^ using deionized water. The Zetasizer (Nano ZS, Malvern, UK) was employed for the comprehensive characterization of PANP, covering particle size, polydispersity index (PDI), and zeta potential. Morphological observations of PANP were conducted by staining the samples with a 2% phosphotungstic acid solution and subsequent examination via transmission electron microscopy (TEM, JEM-2100, JEOL, Japan).

### 2.5. Cell culture

4T1, CT26, B16F10 and NIH/3T3 cell lines were sourced from the Type Culture Collection of the Chinese Academy of Sciences (Beijing, China). LLC cell line was sourced from Dalian Meilun Biotechnology Co., Ltd (Dalian, China). Cell culture media including RPMI 1640, DMEM and DMEM-F12 were enhanced with 10% Fetal Bovine Serum (FBS), streptomycin sulfate, and penicillin. Cellular cultivation took place within a CO_2_-regulated cell incubator at 37°C.

### 2.6. Screening of synergy

First, MTT assay was conducted evaluate the cytotoxicity of co-assembled PANP at various molar ratios. In brief, 4T1 cells were plated in 96-well plates at a density of 1×10^3^ cells per well. After 12 h, media containing varying concentrations of SN38, JQ-1, SN38-LA, JQ-1-LA or co-assembled PANP at various molar ratios were added, followed by 48-h incubation. Post-incubation, MTT solution was introduced, and after 4-h incubation, formazan dissolution was achieved with 200 μL dimethyl sulfoxide (DMSO). The absorbance at 570 nm was recorded by a microplate reader (SYNERGY, BioTek Instruments, Inc, USA). IC_50_ values were calculated on GraphPad Prism 8.0 software. Combination index at IC_50_ (CI_50_) was calculated as the below formula.

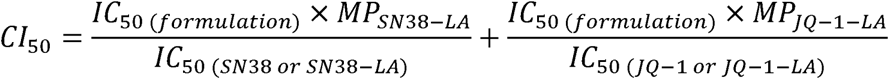

The molar composition (MP) represents the molar proportion of SN38-LA or JQ-1-LA in the formulation. Reference values for IC_50_ were determined using the parent drugs (SN38 and JQ-1) or the respective prodrugs (SN38-LA and JQ-1-LA).

### 2.7. Colloidal stability

The colloidal stability of PANP was assessed through their incubation with PBS (pH 7.4) containing 10% fetal bovine serum (FBS) in a temperature-controlled shaker at 37L for 48 h. Zetasizer measurements were taken at specific time points to monitor changes in particle size. Furthermore, an extended storage experiment was conducted at 4L over a duration of 35 days. Zetasizer measurements were taken during the storage to monitor changes in particle size.

### 2.8. Molecular dynamics simulations

Molecular dynamics (MD) simulations employ a comprehensive model of interatomic interactions to project the dynamic motion of each atom within a protein or molecular system over time [29]. These simulations illuminate essential assembly processes by offering precise atomic positions with femtosecond temporal resolution and insights into intermolecular forces [30]. In our study, the MD simulations were conducted on the Yinfo Cloud Computing Platform (https://cloud.yinfotek.com). To begin with, the 3D conformation of SN38-LA and JQ-1-LA was assessed. Then, the simulations were run by the DOCK 6.9 program. For SNP and JNP, the single SN38-LA or JQ-1-LA, respectively, served as both ligand and receptor molecules. In the case of SJNP, SN38-LA was the ligand, and JQ-1-LA was the receptor. The output conformations were capped at a maximum of 9.

### 2.9. *In vitro* drug release and release mechanisms

The release medium consisting of PBS with 30% ethanol (v/v) was employed in the study. In essence, PANP were combined with the release medium, and this mixture was prepared with or without the addition of H_2_O_2_ in the concentration of 10 mM. Subsequently, the solution was promptly subjected to an oscillator (37°C, 100 r/min). At predetermined time intervals, samples were withdrawn to assess the released drugs. High-Performance Liquid Chromatography (HPLC) was utilized for drug quantification, with SN38 detected at 367 nm, and JQ-1-SH and CA jointly detected at 242 nm. The ROS-responsive drug release mechanisms were mechanistically elucidated by confirming the intermediates formed during the release process using Ultra-Performance Liquid Chromatography-Tandem Mass Spectrometry (UPLC-MS-MS).

### 2.10. Cytotoxicity assay

To evaluate the cytotoxicity of PANP, MTT assays were conducted. In brief, 4T1 and CT26, were plated in 96-well plates at a density of 1×10^3^ cells per well, and B16F10, LLC, and NIH/3T3 cells were plated in 96-well plates at a density of 2× 10^3^ cells per well. After 12 h, media containing varying concentrations of mixed sol (comprising SN38 and JQ-1 at a 1:3 molar ratio) or PANP were added, followed by 48-h incubation. Post-incubation, MTT solution was introduced, and after a 4-h incubation, formazan dissolution was achieved with 200 μL dimethyl sulfoxide (DMSO). The absorbance at 570 nm was recorded by a microplate reader (SYNERGY, BioTek Instruments, Inc, USA). IC_50_ values were calculated on GraphPad Prism 8.0 software. The tumor selective index (SI) was determined as the below formula.

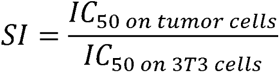

### 2.11. Cellular uptake

In a concise procedure, 4T1 cells were cultured in 24-well plates (5×10^4^ cells per well) with pre-placed lids and allowed to incubate for 24 h. Subsequently, fresh medium containing either free coumarin-6 or coumarin-6-labeled PANP (equivalent to 200 ng mL^−1^ coumarin-6) was introduced, followed by an additional incubation for 0.5 or 2 h. Cold PBS was then used to wash the cells, which were subsequently fixed with a 4% formaldehyde solution. After the fixation process, another round of washing was performed, and Hoechst 33342 staining was applied to visualize the cell nuclei. The cells were finally examined using confocal laser scanning microscopy (CLSM, TCS SP2/AOBS, LEICA, Germany).

### 2.12. Cellular immunofluorescence staining

The immunofluorescence analysis, including γH2AX, BRCA1 and RAD51, was conducted on 4T1 cells post-treatment. Briefly, 4T1 cells were seeded in 24-well plates (10^4^ cells per well) and incubated for 24 h (n = 3 for each group). Subsequently, the cells were exposed to the formulations (200 nM) for a 24-h duration. Blank cells were used as negative control. Following treatment, the cells underwent thrice washing with cold PBS and fixation with 4% formaldehyde. Subsequent to fixation, a distinct antibody (γH2AX, BRCA1 or RAD51) was applied to the cells for an additional 12 h incubation at 4L. Thereafter, the cells underwent thorough washing with cold PBS (five times) to fully remove the antibody. Subsequent to this, cy3-labeled goat anti-rabbit secondary antibody for γH2AX, or FITC-labeled goat anti-rabbit secondary antibody for BRCA1 and RAD51, was administered to the cells for a 1 h duration at room temperature. Following another set of cold PBS washes (five times) to fully remove the secondary antibody, the nuclei were counterstained with Hoechst 33342. The prepared sample slides were observed utilizing CLSM (TCS SP2/AOBS, LEICA, Germany). Quantitative analysis of fluorescence intensity was performed using ImageJ software.

### 2.13. Intracellular ROS level

Intracellular ROS levels were evaluated utilizing the DCFH-DA (ROS fluorescence probe) on 4T1 cells post-treatment. To elaborate, 4T1 cells were cultured on cover glasses and placed in 12-well plates at a density of 10^5^ cells per well. After a 24 h incubation, the cells underwent treatment with the formulations (200 nM) for 48 h. Blank cells were used as negative control. Subsequently, fresh medium containing 10 μM DCFH-DA was introduced and incubated for 30 min at 37°C. The visualization of the cells was then conducted using CLSM (TCS SP2/AOBS, LEICA, Germany)

### 2.14. Michael addition reaction between CA and GSH

The Michael addition reaction between intracellular sulfhydryl compounds and CA was assessed. In brief, CA was diluted to 200 ng mL^−1^ by 30% acetonitrile containing 10 mM glutathione (GSH), and the reaction was kept at 37°C for 30 min. Subsequently, the formation of CA-GSH adducts was verified through UPLC-MS-MS (SHIMADZU CO.,LTD, Kyoto, Japan.)

### 2.15. Intracellular GSH level

To scrutinize alterations in intracellular GSH concentrations post treatment, 4T1 cells were cultured in 6-well plates at a density of 2×10^5^ cells per well. After a 24 h incubation, the cells were treated with the formulations (200 nM) for 12 hours. Subsequently, the cells were harvested and subjected to the instructions of GSH content detection kit procedure. The quantification of GSSG content was recorded by a microplate reader (SYNERGY, BioTek Instruments, Inc, USA).

### 2.16. Mitochondrial permeability transition

Mitochondrial permeability transition was evaluated utilizing the JC-1 probes on 4T1 cells post-treatment. To elaborate, 4T1 cells were cultured on cover glasses and placed in 12-well plates at a density of 1×10^5^ cells per well. After a 24 h incubation, the cells underwent treatment with the formulations (200 nM) for 48 h. CCCP (10 μM) were given to the cells for 20 min as a positive control, and blank cells were used as negative control. Subsequently, fresh medium containing JC-1 reagent (5 μg mL^−1^) was introduced and incubated for 20 min at 37°C in the dark. Finally, the medium was removed, and the cells were washed with JC-1 working buffer for three times. The visualization of the cells was then conducted using CLSM (TCS SP2/AOBS, LEICA, Germany).

### 2.17. *In vivo* Biodistribution

Firstly, we established a subcutaneous 4T1 tumor-bearing mice model. 4T1 cells (5×10^6^ cells) at a volume of 100 μL were subcutaneously inoculated into the dorsal region of female BALB/c mice. When the tumor volume reached approximately 400 mm^3^, we randomly allocated the mice into different groups (n = 3). Subsequently, DiR sol and DiR-labeled PANP were administered intravenously to the mice (equivalent to 1 mg kg^−1^ DiR). The mice were sacrificed at 4 h and 12 h post-administration to dissect the tumors, heart, liver, spleen, lung, and kidney for fluorescence intensity analysis using an IVIS system (IVIS® Lumina III, PerkinElmer, USA).

### 2.18. *In vivo* antitumor performance

To investigate the *in vivo* antitumor performance of the formulations, we established a subcutaneous 4T1 tumor-bearing mice model. 4T1 cells (5×10^6^ cells) at a volume of 100 μL were subcutaneously inoculated into the dorsal region of female BALB/c mice. Upon reaching a size of approximately 100 mm^3^, the mice were randomly allocated into distinct groups (n = 5). Subsequent to the grouping, the mice received PBS or the formulations at doses of 6.4 μM kg^−1^ via the tail vein. Specifically, the dosages included 1.6 μM kg^−1^ SN38 and 4.8 μM kg^−1^ JQ-1 for the mixed sol, 6.4 μM kg^−1^ SN38 for SNP, 6.4 μM kg^−1^ JQ-1 for JNP, and 1.6 μM kg^−1^ SN38 and 4.8 μM kg^−1^ JQ-1 for SJNP. The treatment protocol encompassed five injections administered every other day. Throughout the treatment period, the mice underwent daily measurements of body weight and tumor volume. Upon completion of the study, the tumors were harvested and weighed. Additionally, tumors from each group were dissected for subsequent staining. Blood samples were collected and centrifuged to obtain serum for hepatorenal function analysis using a hepatorenal function kit. Tumor volume and tumor burden were calculated as the below formula.

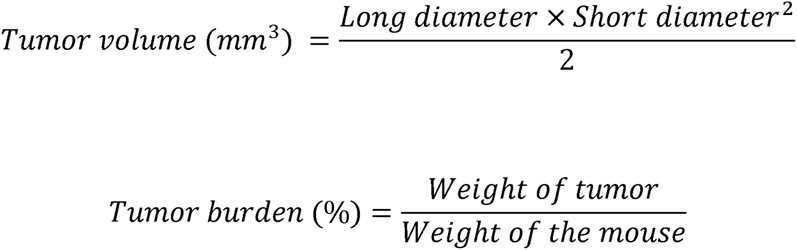

### 2.19. Tumor sections preparation and H&E staining

The excised tumor samples were fixed with 4% paraformaldehyde in PBS and embedded in paraffin. Sections were prepared in 4 μm thick, and subjected to subsequent paraffin clearance and hydration. Hematoxylin solution and eosin Y solution were applied for consecutive staining, and the sections were then dehydrated with ethanol and cleared using xylene, ultimately sealed with Permount. Optical microscopy (Nikon, Tokyo, Japan) was employed for the examination of the H&E prepared samples.

### 2.20. TUNEL assay

Apoptotic cells were discerned through a TUNEL apoptosis assay kit. In brief, following paraffin clearance and hydration, tumor sections underwent pretreatment with Protease K and 0.1% Triton. Subsequently, the sections were incubated with a mixture comprising TDT enzyme, dUTP, and reaction buffer at 37L for 2 hours. Nuclei were stained with DAPI, and the sections were sealed in antifade mounting medium. CLSM (TCS SP2/AOBS, LEICA, Germany) was utilized for the observation of the TUNEL assay.

### 2.21. Immunofluorescence staining of tumor sections

Following paraffin clearance and hydration, tumor sections underwent heat-induced antigen retrieval and serum blocking. Maintained at 4L, the sections were incubated overnight with primary antibodies targeting γH2AX, BRCA1, RAD51, and ki67, individually. Subsequently, a Cy3-labeled secondary antibody was applied at room temperature for 1 hour, followed by nuclear staining utilizing DAPI. The sections were sealed in antifade mounting medium, and observed by CLSM (TCS SP2/AOBS, LEICA, Germany).

### 2.22. Statistical Analysis

Data are portrayed as the mean value ± standard deviation. Statistical analysis was performed using student’s T test (two-tailed) or one-way analysis of variance (ANOVA) to assess the statistical differences between groups. All data were analyzed by GraphPad Prism 8.0. Statistical significance was determined was defined as P < 0.05.

## 3. Results and discussion

### 3.1 Construction of SJNP

In the quest for the proficient co-administration of SN38 and JQ-1, we chose PANP to craft a co-delivery formulation (**Fig. 1**). First, we utilized LA to modify SN38 and JQ-1 (carboxylic acid) through a ROS responsive CT bond to synthesize SN38-LA and JQ-1-LA prodrugs (Fig. S2-3). Both SN38-LA and JQ-1-LA could form PANP with spherical morphology and uniform particle size, namely SNP and JNP (**Fig. 2A** and Table S1). Then, the co-assembly capability of SN38-LA and JQ-1-LA was assessed by preparing PANP at different molar ratios (10:1, 5:1-1:5 and1:10, SN38-LA to JQ-1-LA). At all these ratios, the two prodrugs could co-assemble into PANP, each maintaining a particle size around 80-100 nm (Table. S2).

**Fig. 1.**
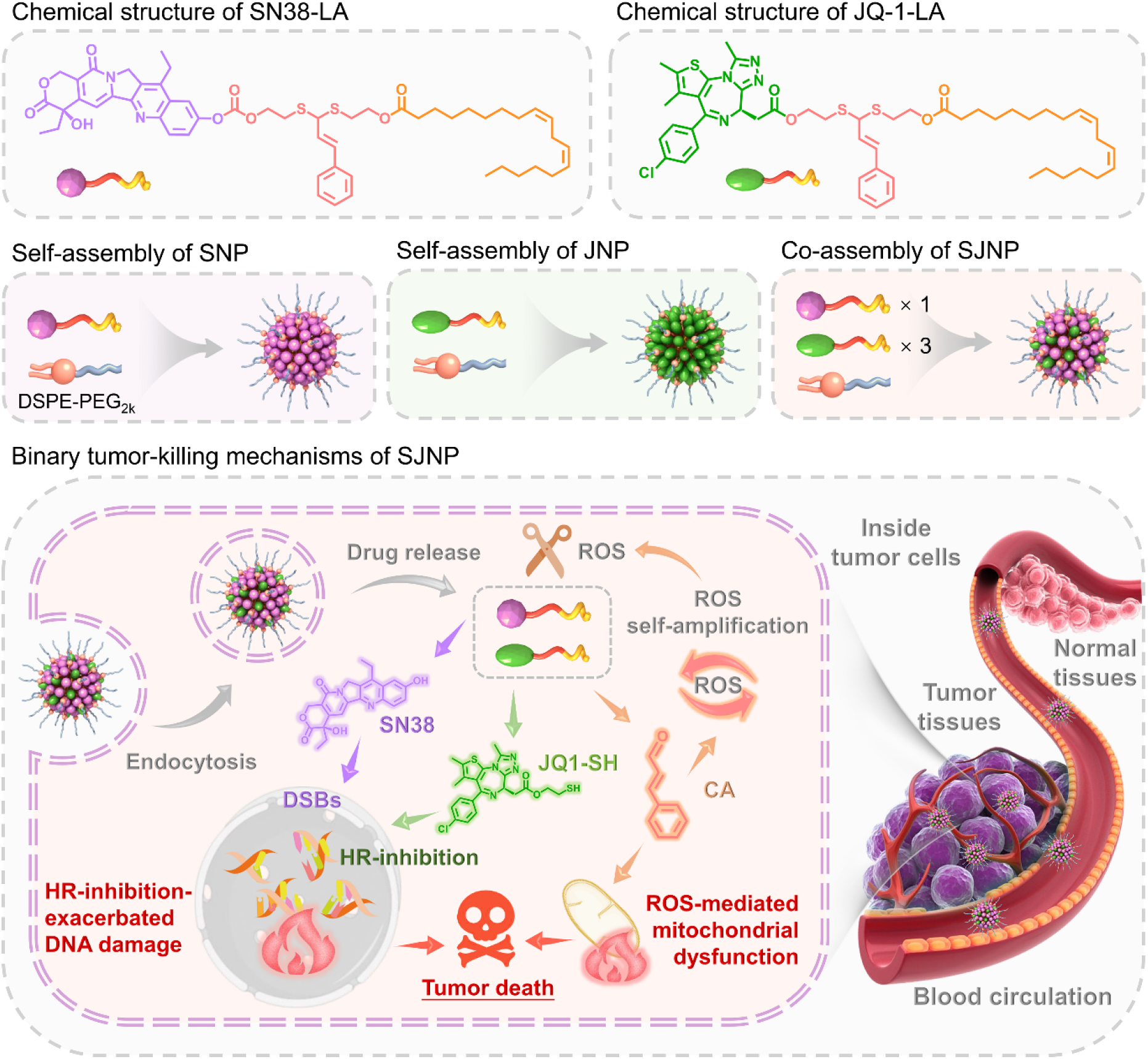
Schematic diagram of the binary tumor-killing mechanisms of prodrug assembled nanoparticles (PANP) for co-delivering SN38 and JQ-1 (SJNP). In the reactive oxygen species (ROS)-enriched tumor microenvironment (TME), SJNP exhibits the capability to liberate active SN38 and JQ-1, along with the by-product cinnamaldehyde (CA). SN38 instigates primary DNA damage, while JQ-1 disrupts BRD4, leading to the downregulation of BRCA1 and RAD51 expression, thereby impairing the homologous recombination (HR) process. The combined effect of SN38 and JQ-1 results in an exacerbation of DNA damage through HR inhibition. Furthermore, CA amplifies cellular ROS levels in a self-reinforcing manner, ultimately inducing ROS-mediated mitochondrial dysfunction. By leveraging these dual mechanisms of tumor eradication, SJNP effectively combats murine triple-negative breast cancer (TNBC) with minimal adverse toxicity.

**Fig. 2.**
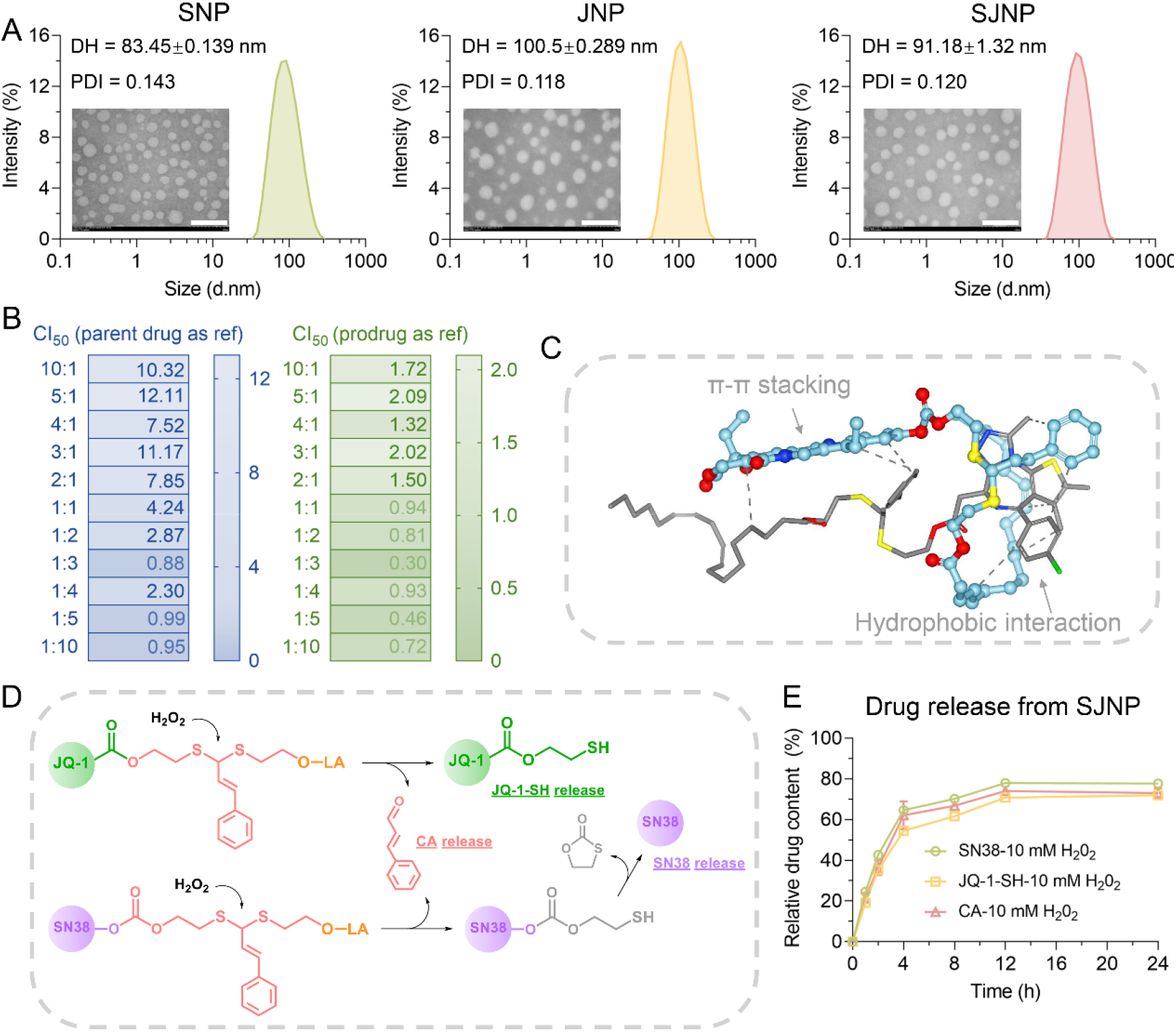
Construction and characterization of PANP. (A) Particle size distribution and transmission electron microscopy (TEM) imaging of PANP. Scale bar = 100 nm. (B) CI_50_ values of co-assembled PANP at different molar ratio. (C) Molecular configuration of SJNP (SN38-LA and JQ-1-LA) during molecular dynamics (MD) simulations. Blue structure represents SN38-LA and gray structure represents JQ-1-LA. (D) Reactive oxygen species (ROS)-responsive release mechanism for SN38-LA and JQ-1-LA. (E) Comparative drug release profiles of SJNP in 10 mM H_2_O_2_. Data are portrayed as mean ± standard deviation (n = 3).

Afterwards, we employed MTT assay to screen the combination index at IC_50_ (CI_50_) of PANP on murine triple-negative breast cancer 4T1 cells. The CI_50_ values were computed using both the parent drug and the prodrug as references, where a CI_50_ < 1 denotes synergy. The outcomes, detailed in Table. S3 and **Fig. 2B**, revealed that at the 1:3 ratio, the CI_50_ reached its minimum (0.88 with the parent drug as a reference and 0.30 with the prodrug as reference), signifying optimal synergy. Consequently, PANP at this 1:3 ratio, designated as SJNP, emerged as the quintessential co-delivery formulation. Notably, SJNP also exhibited spherical morphology and uniform particle size.

### 3.2 Characterization of SJNP

To stimulate the protein-rich *in vivo* component, PANP underwent 48-h incubation in PBS (pH 7.4) supplemented with 10 % FBS at 37°C. This resulted in minimal changes in particle size (Fig. S4A). Further stability was confirmed by a 35-day storage period at 4°C, which showed no significant alterations in particle size (Fig. S4B), demonstrating the robust stability of PANPs suitable for *in vivo* applications and long-term storage.

Molecular dynamics (MD) simulations probed the self-assembly mechanisms of PANPs. For SNP, aromatic π-π stacking and hydrophobic interactions between LA chains were observed (Fig. S5A). In contrast, the self-assembly of JNP was primarily driven by hydrophobic interactions (Fig. S5B). In the co-assembly of SJNP, the aromatic components of SN38 exhibited π-π stacking with the aromatic ring of the CT linkage in JQ-1-LA (**Fig. 2C**). The LA chains from SN38-LA engaged with the JQ-1 structure through shared hydrophobic interactions, culminating in an efficient co-assembly process.

Subsequently, the ROS-responsive drug release profiles were investigated to elucidate the mechanisms (**Fig. 2D**). UPLC-MS-MS analysis confirmed the release intermediates (Fig. S6). ROS exposure cleaved the CT bond, releasing cinnamaldehyde (CA) and generating carbonate-bridged SN38-SH and ester-bridged JQ-1-SH intermediates. The SN38-SH thiol group initiated a nucleophilic attack on the carbonate bond, accelerating a self-elimination cyclization to release SN38. Meanwhile, JQ-1-SH underwent hydrolysis of its ester bond to release JQ-1, with both sharing equivalent pharmacological effects.

The in vitro ROS-responsive release profiles were characterized by the accumulated concentration of CA, SN38, and JQ-1-SH. In the presence of 10 mM H_2_O_2_, mimicking ROS conditions, SJNP released approximately 80% of CA and the respective parent drugs within 24 hours. In contrast, less than 10% of the drugs were released in PBS alone (**Fig. 2E** and Fig. S7). This pronounced difference underscores the distinctive ROS-responsiveness of PANPs, highlighting their ability to differentiate between ROS-overexpressing tumor cells and normal cells.

### 3.3 BRD4 inhibition exacerbates DSBs by impairing HR

We commenced by assessing the cytotoxic effects of our formulations across a spectrum of tumor cell lines, including 4T1, CT26, B16F10, and LLC. Analysis revealed that, while the NP formulation showed a slight reduction in cytotoxicity compared to the mixed drug solution (sol), this was likely due to the controlled release characteristics of the prodrugs (**Fig. 3A** and Fig. S8). Notably, the SJNP formulation displayed significantly increased cytotoxicity relative to SNP and JNP, underscoring the enhanced antiproliferative potential of the combined approach. Additionally, we evaluated the cytotoxicity against normal NIH/3T3 cells (Fig. S9) and calculated the selectivity index (SI) (Table S4). The elevated SI observed for SJNP suggests a pronounced tumor selectivity, which may contribute to a better safety profile.

**Fig. 3.**
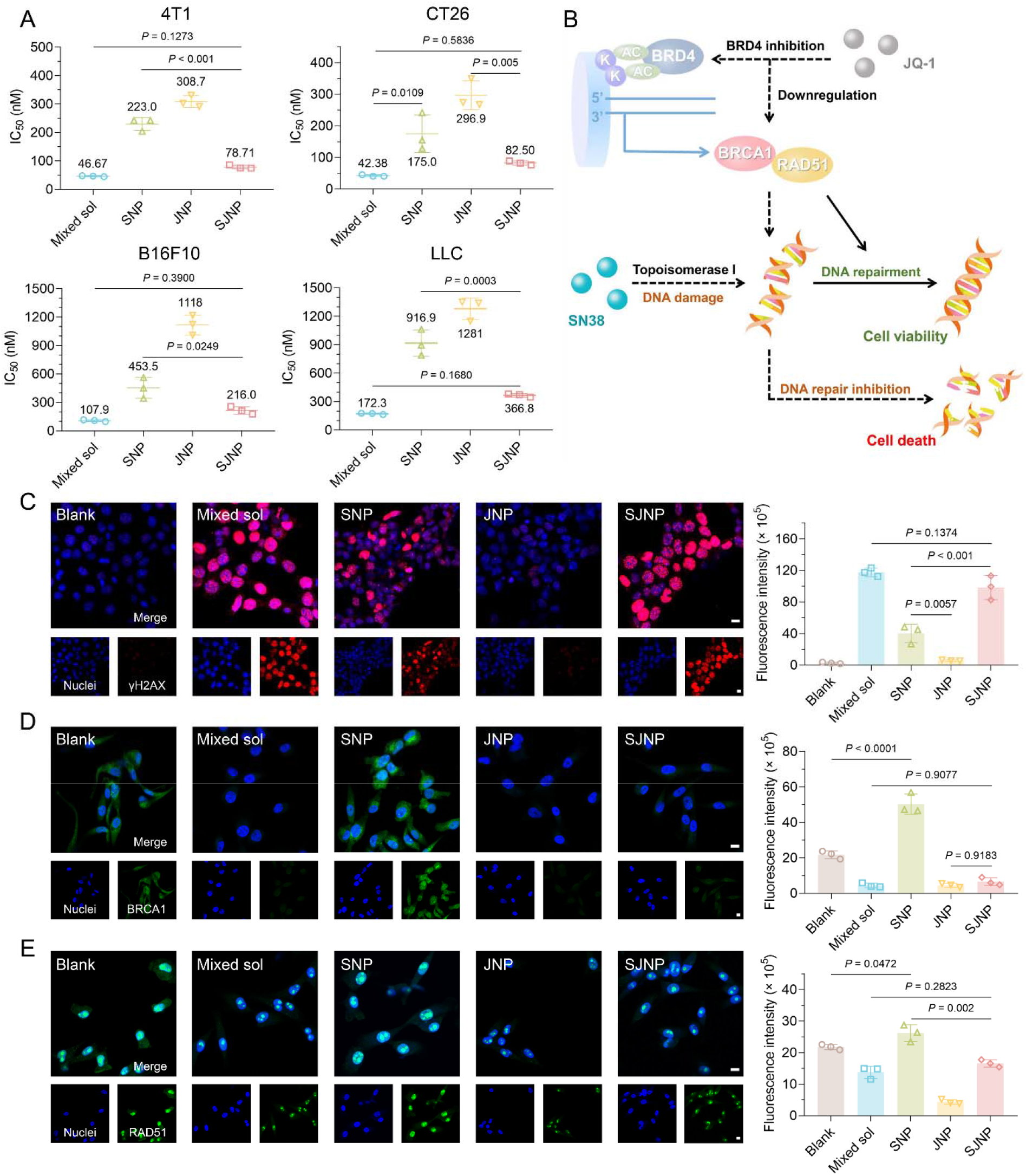
BRD4 inhibition exacerbates DSBs by impairing HR. A) Cytotoxicity of PANP on 4T1, CT26, B16F10 and LCC cells. Data are portrayed as mean ± standard deviation (n = 3). Statistical significance was assessed through ANOVA and deemed significant at P < 0.05. B) Proposed mechanisms of BRD4 inhibition exacerbates DSBs by impairing HR. Dashed lines guide tumor cell death, while solid lines may lead to tumor survival. (C, D, and E) Immunofluorescence and quantitative results of C) γH2AX, D) BRCA1 and E) RAD51. Scale bar = 10 μm. Data are portrayed as mean ± standard deviation (n = 3). Statistical significance was assessed through ANOVA and deemed significant at P < 0.05.

The efficacy of PANP against cancer cells appears tightly linked to their uptake by these cells. We quantified cellular internalization by tracking the fluorescence intensity of coumarin-6-labeled PANP in 4T1 cells. Data indicated that PANP internalization was significantly higher than that of free coumarin-6 and increased progressively over time (Fig. S10). Notably, SNP, JNP, and SJNP showed similar uptake efficiencies, likely due to comparable surface properties such as particle size and zeta potential. These results support the premise that differences in cytotoxicity among PANP formulations are primarily due to their distinct mechanisms of action, rather than differences in cellular uptake.

To elucidate the molecular dynamics of the proposed synergetic effect of SN38 and JQ-1 (**Fig. 3B**), we performed immunofluorescence studies on 4T1 cells treated with our formulations at a total drug concentration of 200 nM. The assay focused on γH2AX expression, a marker for double-strand breaks (DSBs). The results indicated that the synergistic formulation, both the mixed solution and SJNP, induced substantial γH2AX expression, with SNP also showing a notable increase. By contrast, JNP, which lacks the ability to induce DNA damage, exhibited minimal γH2AX expression, correlating with its lower cytotoxic impact (**Fig. 3C**).

Double-strand breaks activate the HR repair pathway, where the transcription factor BRD4 modulates critical HR components, including BRCA1 and RAD51. The SNP-induced DNA damage led to a marked upregulation of BRCA1 and RAD51, suggesting an enhanced HR response (**Fig. 3D** and **E**). Conversely, JNP-mediated BRD4 inhibition resulted in a marked decrease in the expression of these factors compared to controls. The inclusion of JQ-1 in SJNP contributed to the downregulation of BRCA1 and RAD51, mirroring the effect observed with the mixed solution. Consequently, the combined effect in SJNP compromised HR, thereby exacerbating DSBs. This mechanistic insight aligns with the observation that SJNP elicits a more potent DNA damage response and increased γH2AX expression, even at a lower SN38 concentration than SNP.

### 3.4 ROS self-amplification culminates in mitochondrial dysfunction

In the intricate design of PANP, besides serving as a chemical linkage, the CT structure could also self-deliver another antitumor component CA. Previous studies have shown that CA and its derivatives can suppress the growth of various cancer cells by promoting the production of intracellular ROS. Our results corroborate these findings, with PANP-treated 4T1 cells displaying a substantial rise in intracellular ROS compared to cells treated with the mixed drug solution (sol) or left untreated (**Fig. 4A**). The α, β-unsaturated carbonyl group in CA, recognized as a potent Michael acceptor, likely contributes to this ROS increase by reacting with glutathione (GSH) through a Michael addition reaction. This reaction reduces intracellular GSH levels, subsequently enhancing ROS concentration (**Fig. 4B**). Quantitative analysis of GSH levels within 4T1 cells further revealed a significant depletion of GSH in PANP-treated cells, an effect not observed with the mixed sol (**Fig. 4C**). Besides, the formation of CA-GSH adducts using UPLC-MS-MS analysis (**Fig. 4D**).

**Fig. 4.**
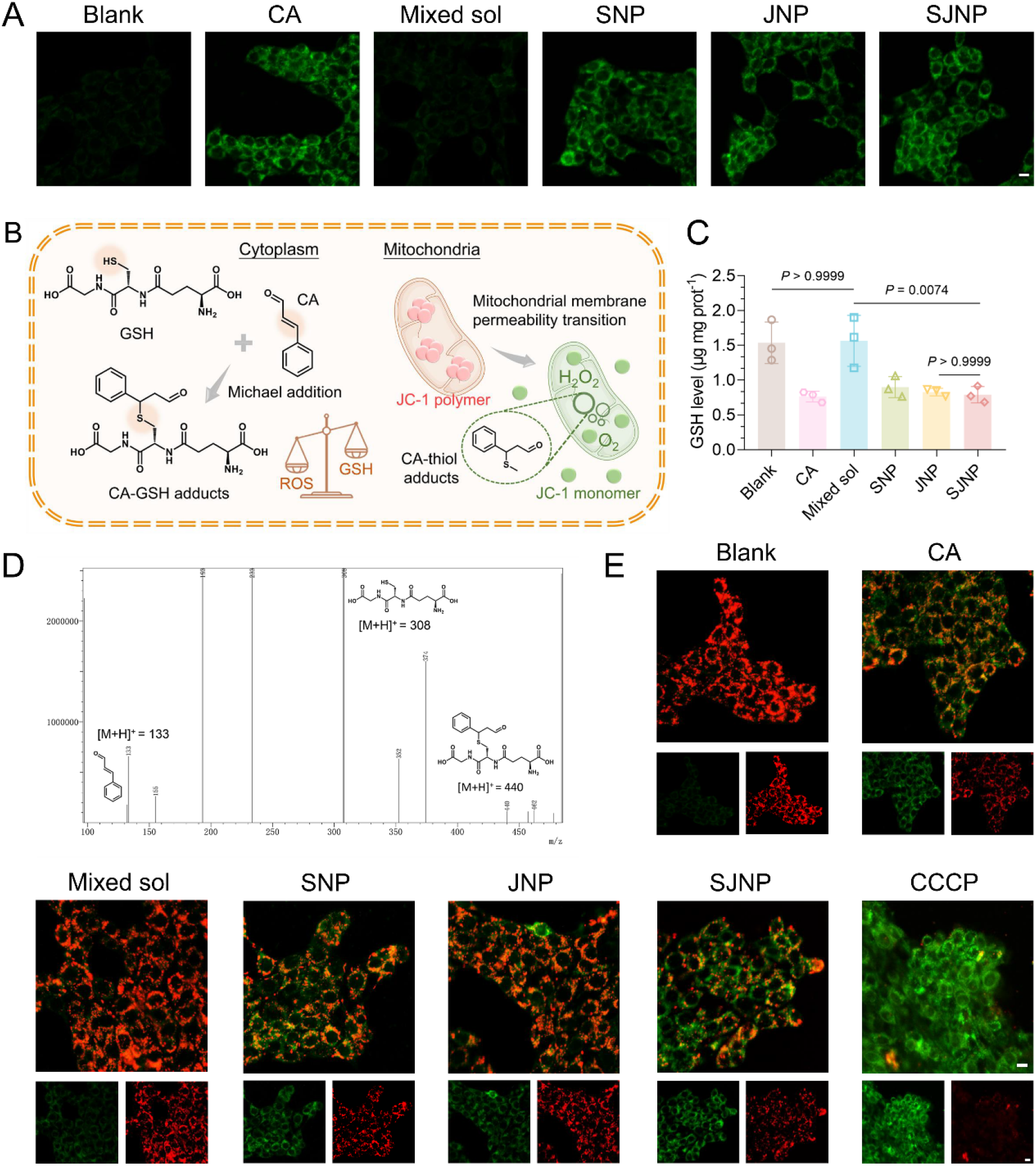
ROS self-amplification culminates mitochondrial dysfunction. A) Intracellular ROS level detected by DCFH-DA probe of 4T1 cells post administration. Scale bar = 10 μm. B) Schematic illustration of intracellular formation of CA-GSH adducts, and mitochondrial membrane permeability transition. C) Cellular GSH concentration of 4T1 cells post administration. Data are portrayed as mean ± standard deviation (n = 3). Statistical significance was assessed through ANOVA and deemed significant at P < 0.05. D) Structure conformation of the formation of CA-GSH adducts by UPLC-MS-MS. E) Mitochondrial permeability transition detected by JC-1 probe of 4T1 cells post administration. Carbonyl cyanide 3-chlorophenylhydrazone (CCCP) is employed as positive control. Scale bar = 10 μm.

Within tumor cells, the CA released in response to elevated ROS levels can further augment intracellular ROS, creating a self-perpetuating cycle of ‘ROS self-amplification.’ This novel mechanism overcomes a key hurdle in prodrug-based therapies—efficient bioactivation—thereby setting the stage for the activated drugs to exert their anticancer effects more effectively.

Additionally, CA’s interaction with mitochondrial thiol-containing antioxidants disrupts the mitochondrial redox balance, potentially leading to ROS-induced mitochondrial permeability transition and subsequent mitochondrial dysfunction. We investigated this by examining mitochondrial membrane permeability using the JC-1 dye, where functional mitochondria exhibit red fluorescence due to JC-1 aggregation, and dysfunctional mitochondria exhibit green fluorescence due to JC-1 monomer dispersal (**Fig. 4A**). PANP-treated cells, and notably those treated with SJNP, showed an increased green fluorescence, indicative of mitochondrial dysfunction (**Fig. 4E**). Despite the absence of CA in the mixed sol, we observed a slight green fluorescence, attributed to apoptosis-induced mitochondrial permeability transition, which was more pronounced in cells treated with SJNP as compared to SNP and JNP.

In summary, the ROS self-amplification characteristic of our PANP design not only ensures efficient tumor-responsive drug release but also contributes to the therapeutic efficacy. The remarkable anticancer activity of SJNP can be attributed to the combined effects of HR inhibition, which intensifies DNA damage, and ROS-mediated mitochondrial dysfunction.

### 3.5 Antitumor efficacy of SJNP in a Triple-Negative Breast Cancer model

Ensuring the efficient accumulation at tumor sites is pivotal for unlocking the antitumor potential of PANP. Prior to the *in vivo* antitumor evaluation, we first explored the biodistribution of DiR-labeled PANP in a 4T1 murine TNBC model. Imaging results revealed the rapid distribution of both free DiR and DiR-labeled PANP across major organs (**Fig. 5A** and S11). Crucially, within the tumor tissues, PANP showed superior accumulation over free DiR, with signal intensity peaking at 12 hours post-administration, suggestive of successful tumor homing via the EPR effect (**Fig. 5B**). Importantly, no significant differences were observed among the various PANP formulations, likely due to their similar surface characteristics affecting their *in vivo* behavior.

**Fig. 5.**
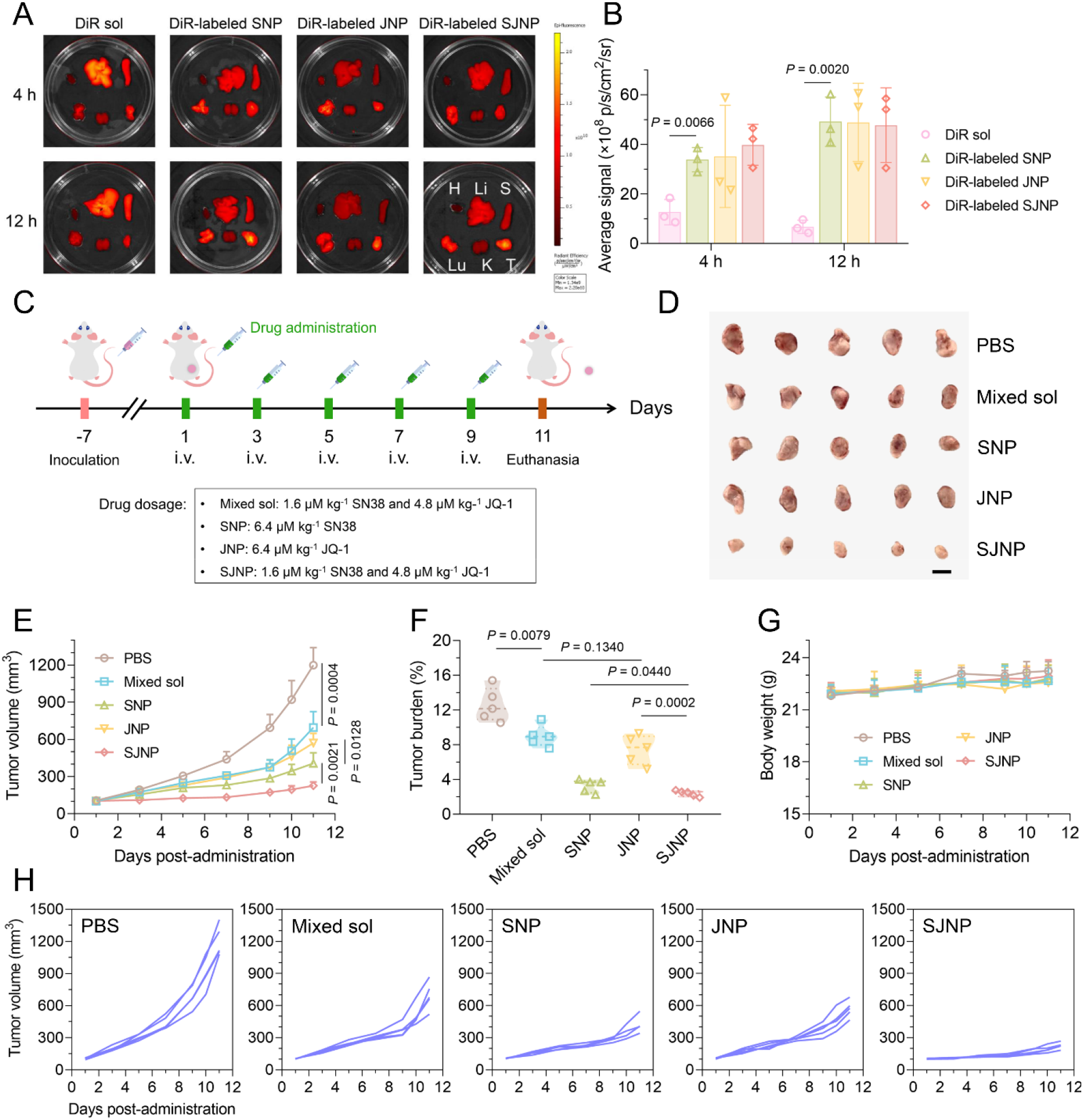
SJNP significantly suppresses murine TNBC growth. A) Biodistribution of DiR-labeled PANP. H: heart; Li: Liver; S: spleen; Lu: lung; K: kidney; T: tumor. B) Quantitative results of tumor accumulation of DiR-labeled PANP. Data are portrayed as mean ± standard deviation (n = 3). Statistical significance was assessed through ANOVA and deemed significant at P < 0.05. C) Protocol of pharmacodynamics study. D) Images of dissected 4T1 tumors at the end of treatment. Scale bar = 1 cm. E) Tumor volume curves during the treatment. Data are portrayed as mean ± standard deviation (n = 5). Statistical significance was assessed through student’s T test (two-tailed) and deemed significant at P < 0.05. F) Tumor burdens at the end of treatment. Data are portrayed as mean ± standard deviation (n = 5). Statistical significance was assessed through student’s T test (two-tailed) and deemed significant at P < 0.05. G) Body weight curves during the treatment. H) Individual tumor growth curves of the mice during the treatment period.

We then assessed the in vivo antitumor efficacy using the 4T1 TNBC model. The pharmacodynamic study protocol is detailed in **Fig. 5C**. Tumor-bearing mice were treated intravenously with the formulations at a dose of 6.4 μM kg^−1^ every other day, for a total of five doses. Tumor progression and systemic toxicity were monitored by measuring tumor volume and body weight, respectively. The antitumor activity was quantified by evaluating the changes in tumor volume and overall tumor burden (**Fig. 5D-F** and **H**). The control group, treated with PBS, showed rapid tumor progression. In comparison, the mixed sol and JNP groups exhibited modest tumor growth inhibition. In contrast, SJNP displayed the most substantial inhibition of tumor growth, outperforming SNP. The safety profile of these treatments was corroborated by the stable body weight of the mice and the absence of any detectable hepatorenal toxicity post-treatment (**Fig. 5G** and **Fig. 6A**). These data highlight the superior antitumor performance of SJNP and confirm its safety, presenting no additional toxicity concerns.

**Fig. 6.**
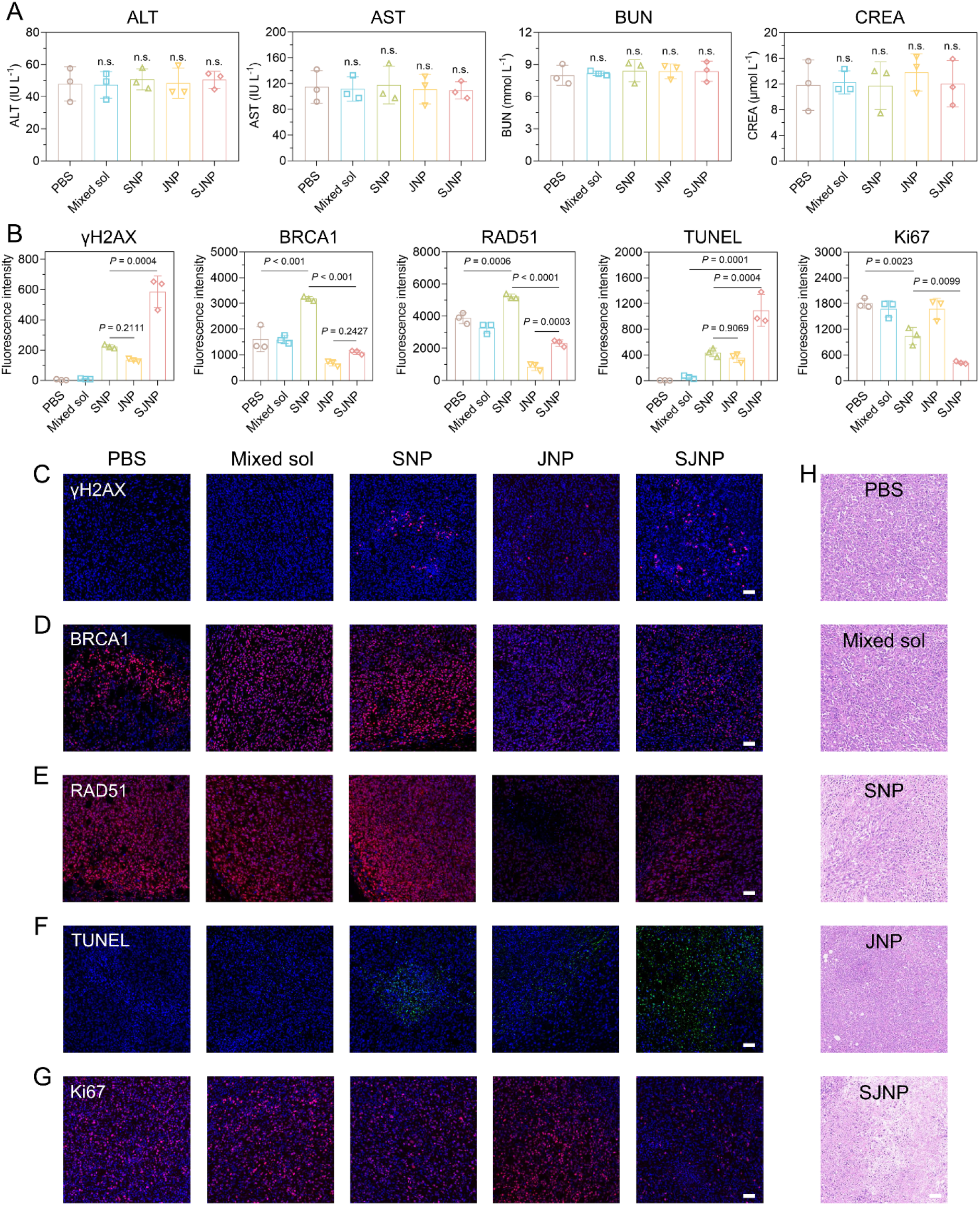
SJNP grantees biosafety of treatment without additional hepatorenal toxicity, and elicits binary tumor-killing mechanisms on murine TNBC model. A) Hepatorenal function parameters of of the mice at the end of treatment. Data are portrayed as mean ± standard deviation (n = 3). Statistical significance was assessed through ANOVA and deemed unsignificant at P > 0.05. B) Quantitative results of TUNEL assay and immunofluorescence staining. Data are portrayed as mean ± standard deviation (n = 3). Statistical significance was assessed through ANOVA and deemed significant at P < 0.05. (C, D, E, F and G) Immunofluorescence staining of C) γH2AX, D) BRCA1, E) RAD51, F) TUNEL assay and G) Ki67. Scale bar = 50 μm. H) H&E staining. Scale bar = 50 μm.

Post-treatment, tumor specimens were procured for detailed histological analysis. Consistent with the extent of necrosis, immunofluorescence assays demonstrated that the SJNP group exhibited the highest levels of γH2AX, a marker for DNA double-strand breaks, surpassing those seen in the SNP group (**Fig. 6B-C**). This was complemented by changes in the expression of BRCA1 and RAD51, key proteins in HR repair. The SNP group showed increased expression of these proteins relative to the control (PBS group), while JNP and the synergistic formulations (mixed sol and SJNP) downregulated their expression (**Fig. 6B** and **D-E**). These findings corroborate the cellular studies, reinforcing the conclusion that SJNP impedes HR and amplifies DNA damage within tumor cells.

Further, terminal deoxynucleotidyl transferase dUTP nick end labeling (TUNEL) assays highlighted that SJNP induced the most extensive apoptosis among tumor cells, followed by SNP, the mixed solution, and JNP, in descending order of effect (**Fig. 6B** and **F**). This trend aligns with the observed impact of ROS-mediated mitochondrial dysfunction in triggering cell death. Conversely, Ki67 staining, a proxy for cell proliferation, displayed a reverse pattern, aligning with the anticancer effects of each treatment (**Fig. 6B** and **G**). Consistent with the extent of TUNEL and Ki67 assay, hematoxylin and eosin (H&E) staining revealed pronounced necrosis within tumors from the SJNP-treated group, indicating significant therapeutic impact (**Fig. 6H**).

The exceptional antitumor activity of SJNP can be attributed to the dual mechanisms of action: the exacerbation of DNA damage through HR inhibition and the induction of mitochondrial dysfunction via ROS generation. Collectively, our study positions SJNP as a strategically engineered nanoplatform, exhibiting robust antitumor efficacy coupled with minimal toxicity.

## 4. Conclusion

The strategic induction of DNA damage is a cornerstone of cancer therapy, exemplified by the clinical deployment of chemotherapeutic agents such as alkylating agents and topoisomerase inhibitors, along with photodynamic therapy and radiotherapy. However, the therapeutic efficacy of DNA damage-inducing strategies is often compromised by cancer cells’ adaptive resistance mechanisms, particularly through modified DNA repair pathways. A concurrent blockade of DNA damage repair mechanisms during DNA damage induction has emerged as a promising strategy to significantly enhance the efficacy of DNA damage-based therapies.

SN38, a topoisomerase I (TOP1) inhibitor, induces cytotoxicity by triggering DSBs during DNA replication and transcription. In response to SN38-induced DSBs, tumor cells typically engage HR repair to restore DNA integrity. BRD4, a transcription factor protein from the BET family, regulates the expression of critical HR components such as BRCA1 and RAD51. To disrupt HR, we have co-opted JQ-1, a BET inhibitor, to synergistically augment SN38’s efficacy. This combination seeks to dismantle the HR repair apparatus while sensitizing tumor cells to SN38-induced DNA damage, offering a novel and targeted therapeutic avenue.

Our research opens pathways for investigating the susceptibility of various tumor subtypes to DNA damage therapies coupled with DNA repair inhibition. Particularly in TNBC, where 20-25% of patients exhibit germline BRCA1/2 mutations or a BRCAness phenotype [31, 32]— characterized by impaired RAD51 complex function or epigenetic modifications affecting BRCA1 expression [33, 34]—our strategy shows potential. However, the effectiveness of our synergistic approach in cases exhibiting the BRCAness phenotype remains to be determined.

Furthermore, the clinical translation of our nanomedicine, SJNP, is not without challenges, such as industrial-scale sterilization, lyophilization, and physiological variances between animal models and human patients.

In this study, we designed PANP to deliver SN38 and JQ-1, incorporating a ROS self-amplifying CT linkage within their prodrug framework. Our comprehensive investigations revealed several therapeutic benefits of SJNP, including a tunable drug loading ratio, tumor-targeted ROS-responsive drug release, and intracellular ROS self-amplification, alongside enhanced cellular uptake, efficient tumor targeting via the EPR effect, and formidable in vivo antitumor efficacy with high biocompatibility.

In both cellular and in vivo TNBC models, SJNP’s dual tumor-killing mechanism was demonstrated. It downregulated BRCA1 and RAD51 to intensify SN38-induced DSBs by disrupting HR and utilized the ROS-responsive cleavage of the CT linkage to release CA, thereby propelling ROS-mediated mitochondrial dysfunction and apoptosis. Collectively, SJNP stands out as a promising nanoplatform, offering high clinical promise and translational potential, and provides a significant leap forward in enhancing DNA damage-based therapies and developing next-generation nanomedicines.

## Supporting information

Methods for chemical synthesis; Supplemental Figure S1-S11; Supplemental Table S1-S4

## Data availability

All relevant data are available from the authors.

## Acknowledgement

This work was financially supported by Liaoning Provincial Department of Education program (LJKMZ20221353), Natural Science Foundation of Liaoning Province (2022-BS-158), National Natural Science Foundation of China (82104109) and the Japan Society for the Promotion of Science (JSPS; 21H01728). The WPI-iCeMS is supported by the World Premier International Research Centre Initiative (WPI), MEXT, Japan.

## Author contributions

Shunzhe Zheng designed the project and wrote this paper; Meng Li analyzed data; Wenqian Xu and Jiaxin Zhang drew the figures; Guanting Li drew the graphic abstract; Hongying Xiao and Xinying Liu collected literatures; Jianbing Shi and Fengli Xia drew the tables, Chuntong Tian* and Ken-ichiro Kamei* offered guidance and funds for this work.

## Reference

[1] J. Ferlay, M. Colombet, I. Soerjomataram, D.M. Parkin, M. Piñeros, A. Znaor, F. Bray, Cancer statistics for the year 2020: An overview, International journal of cancer, 149 (2021) 778–789.

[2] R.L. Siegel, K.D. Miller, N.S. Wagle, A. Jemal, Cancer statistics, 2023, Ca Cancer J Clin, 73 (2023) 17–48.

[3] P. Bouwman, J. Jonkers, The effects of deregulated DNA damage signalling on cancer chemotherapy response and resistance, Nature Reviews Cancer, 12 (2012) 587–598.

[4] L.H. Swift, R.M. Golsteyn, Genotoxic anti-cancer agents and their relationship to DNA damage, mitosis, and checkpoint adaptation in proliferating cancer cells, International journal of molecular sciences, 15 (2014) 3403–3431.

[5] C. Bailly, Irinotecan: 25 years of cancer treatment, Pharmacological research, 148 (2019) 104398.

[6] M. Ramesh, P. Ahlawat, N.R. Srinivas, Irinotecan and its active metabolite, SNL38: review of bioanalytical methods and recent update from clinical pharmacology perspectives, Biomedical chromatography, 24 (2010) 104–123.

[7] M. Kciuk, B. Marciniak, R. Kontek, Irinotecan—still an important player in cancer chemotherapy: a comprehensive overview, International journal of molecular sciences, 21 (2020) 4919.

[8] X. Li, W.-D. Heyer, Homologous recombination in DNA repair and DNA damage tolerance, Cell research, 18 (2008) 99–113.

[9] T. Helleday, Homologous recombination in cancer development, treatment and development of drug resistance, Carcinogenesis, 31 (2010) 955–960.

[10] F.J. Groelly, M. Fawkes, R.A. Dagg, A.N. Blackford, M. Tarsounas, Targeting DNA damage response pathways in cancer, Nature Reviews Cancer, 23 (2023) 78–94.

[11] A.C. Belkina, G.V. Denis, BET domain co-regulators in obesity, inflammation and cancer, Nature reviews Cancer, 12 (2012) 465–477.

[12] N. Wang, R. Wu, D. Tang, R. Kang, The BET family in immunity and disease, Signal transduction and targeted therapy, 6 (2021) 23.

[13] X. Han, D. Yu, R. Gu, Y. Jia, Q. Wang, A. Jaganathan, X. Yang, M. Yu, N. Babault, C. Zhao, Roles of the BRD4 short isoform in phase separation and active gene transcription, Nature Structural & Molecular Biology, 27 (2020) 333–341.

[14] C. Mio, L. Gerratana, M. Bolis, F. Caponnetto, A. Zanello, M. Barbina, C. Di Loreto, E. Garattini, G. Damante, F. Puglisi, BET proteins regulate homologous recombinationLmediated DNA repair: BRCAness and implications for cancer therapy, International journal of cancer, 144 (2019) 755–766.

[15] R. Prakash, Y. Zhang, W. Feng, M. Jasin, Homologous recombination and human health: the roles of BRCA1, BRCA2, and associated proteins, Cold Spring Harbor perspectives in biology, 7 (2015) a016600.

[16] G. Matos-Rodrigues, J. Guirouilh-Barbat, E. Martini, B.S. Lopez, Homologous recombination, cancer and the ‘RAD51 paradox’, NAR cancer, 3 (2021) zcab016.

[17] G. Li, B. Sun, Y. Li, C. Luo, Z. He, J. Sun, SmallLmolecule prodrug nanoassemblies: an emerging nanoplatform for anticancer drug delivery, Small, 17 (2021) 2101460.

[18] S. Zheng, G. Li, J. Shi, X. Liu, M. Li, Z. He, C. Tian, K.-i. Kamei, Emerging platinum (IV) prodrug nanotherapeutics: A new epoch for platinum-based cancer therapy, Journal of Controlled Release, 361 (2023) 819–846.

[19] S. Fu, G. Li, W. Zang, X. Zhou, K. Shi, Y. Zhai, Pure drug nano-assemblies: A facile carrier-free nanoplatform for efficient cancer therapy, Acta Pharmaceutica Sinica B, 12 (2022) 92–106.

[20] G. Li, B. Sun, S. Zheng, L. Xu, W. Tao, D. Zhao, J. Yu, S. Fu, X. Zhang, H. Zhang, ZwitterionLdriven shape program of prodrug nanoassemblies with high stability, high tumor accumulation, and high antitumor activity, Advanced Healthcare Materials, 10 (2021) 2101407.

[21] G. Li, F. Xia, H. Xiao, S. Zheng, S. Fu, H. Qiao, Q. Jin, X. Zhang, D. Zhou, C. Tian, Fine-tuning the structure-tolerance-antitumor efficacy axis of prodrug nanoassemblies via branched aliphatic functionalization, Nano Research, (2023) 1–11.

[22] G. Li, Q. Jin, F. Xia, S. Fu, X. Zhang, H. Xiao, C. Tian, Q. Lv, J. Sun, Z. He, Smart stimuli-responsive carrier-free nanoassembly of SN38 prodrug as efficient chemotherapeutic nanomedicine, Acta Materia Medica, 2 (2023) 54–63.

[23] S. Zheng, G. Li, S. Fu, N. Wang, H. Qiao, M. Li, X. Zhang, K. Wang, W. Sun, C. Tian, Hybrid nanoassembly indicating a synthetic lethality relationship induces mitotic catastrophe-mediated tumor elimination, Chemical Engineering Journal, 479 (2024) 147802.

[24] P. Jangili, N. Kong, J.H. Kim, J. Zhou, H. Liu, X. Zhang, W. Tao, J.S. Kim, DNALdamageLresponseLtargeting mitochondriaLactivated multifunctional prodrug strategy for selfLdefensive tumor therapy, Angewandte Chemie International Edition, 61 (2022) e202117075.

[25] M. Chen, D. Liu, F. Liu, Y. Wu, X. Peng, F. Song, Recent advances of redox-responsive nanoplatforms for tumor theranostics, Journal of Controlled Release, 332 (2021) 269–284.

[26] F. Gong, N. Yang, X. Wang, Q. Zhao, Q. Chen, Z. Liu, L. Cheng, Tumor microenvironment-responsive intelligent nanoplatforms for cancer theranostics, Nano Today, 32 (2020) 100851.

[27] L. Chaiswing, W.H. St. Clair, D.K. St. Clair, Redox paradox: a novel approach to therapeutics-resistant cancer, Antioxidants & redox signaling, 29 (2018) 1237–1272.

[28] S.S. Sabharwal, P.T. Schumacker, Mitochondrial ROS in cancer: initiators, amplifiers or an Achilles’ heel?, Nature Reviews Cancer, 14 (2014) 709–721.

[29] M. Karplus, J.A. McCammon, Molecular dynamics simulations of biomolecules, Nature structural biology, 9 (2002) 646–652.

[30] S.A. Hollingsworth, R.O. Dror, Molecular dynamics simulation for all, Neuron, 99 (2018) 1129–1143.

[31] J.H. Park, J.-H. Ahn, S.-B. Kim, How shall we treat early triple-negative breast cancer (TNBC): from the current standard to upcoming immuno-molecular strategies, ESMO open, 3 (2018) e000357.

[32] I. Gorodetska, I. Kozeretska, A. Dubrovska, BRCA genes: the role in genome stability, cancer stemness and therapy resistance, Journal of Cancer, 10 (2019) 2109.

[33] C.J. Lord, A. Ashworth, BRCAness revisited, Nature Reviews Cancer, 16 (2016) 110–120.

[34] M. Moschetta, A. George, S. Kaye, S. Banerjee, BRCA somatic mutations and epigenetic BRCA modifications in serous ovarian cancer, Annals of Oncology, 27 (2016) 1449–1455.

