## Supplementary material for "Dual-targeted nanoparticulate drug delivery systems for enhancing triple-negative breast cancer treatment": Methods for chemical synthesis; Supplemental Figure S1-S11; Supplemental Table S1-S4

The chemical synthesis route was illustrated in Fig. S1.

**1. Synthesis of CT**

In a glass flask, mercaptoethanol (5 mmol), CA (2 mmol), and hydrochloric acid (10 mmol) were combined and stirred at 0 °C for 30 min under an argon atmosphere. The resulting viscous solution was washed with water to eliminate the majority of hydrochloric acid. The purification process involved column chromatography to yield the CT linkage. The product was light yellow solid (yield: 76.4%).

**2. Synthesis of LA-CT**

In methylbenzene, LA (0.8 mmol) and CT (1.5 mmol) were dissolved, followed by the addition of p-toluenesulfonic acid (1.2 mmol). The reaction was maintained at 110℃ under reflux for 2 h with vigorous stirring. The reaction completion was verified through silica gel thin-layer chromatography (TLC). Following three washes with NaHCO_3_, the product underwent purification via column chromatography to yield LA-CT, an oily liquid with a yield of 63.3%.

**3. Synthesis of SN38-LA**

In anhydrous dichloromethane (DCM), LA-CT (0.4 mmol) underwent dissolution, with the addition of N,N-diisopropylethylamine (0.8 mmol) and p-nitrophenyl chloroformate (0.4 mmol). The reaction persisted at 25℃ for 12 h under vigorous stirring. Subsequently, DCM was eliminated through rotary evaporation. Without additional purification, the crude product was redissolved in anhydrous N,N-dimethylformamide (DMF). Triethylamine (0.8 mmol) and SN38 (0.4 mmol) were introduced into the solution, maintaining the reaction at 25℃ for 12 h with vigorous stirring. Eventually, the product was extracted into ethyl acetate. The resulting compound underwent purification using preparative liquid chromatography to yield SN38-LA (yield: 71.3%). Structural confirmation was achieved through mass spectrometry (MS) and nuclear magnetic resonance spectroscopy (NMR).

**4. Synthesis of JQ-1-LA**

In anhydrous dichloromethane, a solution was formed by dissolving JQ-1 (carboxylic acid) (0.4 mmol), LA-CT (0.5 mmol), 1-ethyl-3(3-dimethylpropylamine) carbodiimide (0.6 mmol), 4-dimethylaminopyridine (0.15 mmol), and 1-hydroxybenzotriazole (0.6 mmol). The reaction mixture underwent stirring at 25 °C for 24 h. Eventually, the product was extracted into ethyl acetate. The resulting compound underwent purification using preparative liquid chromatography to yield JQ-1-LA (yield: 55.4%). Structural confirmation was achieved through MS and NMR.

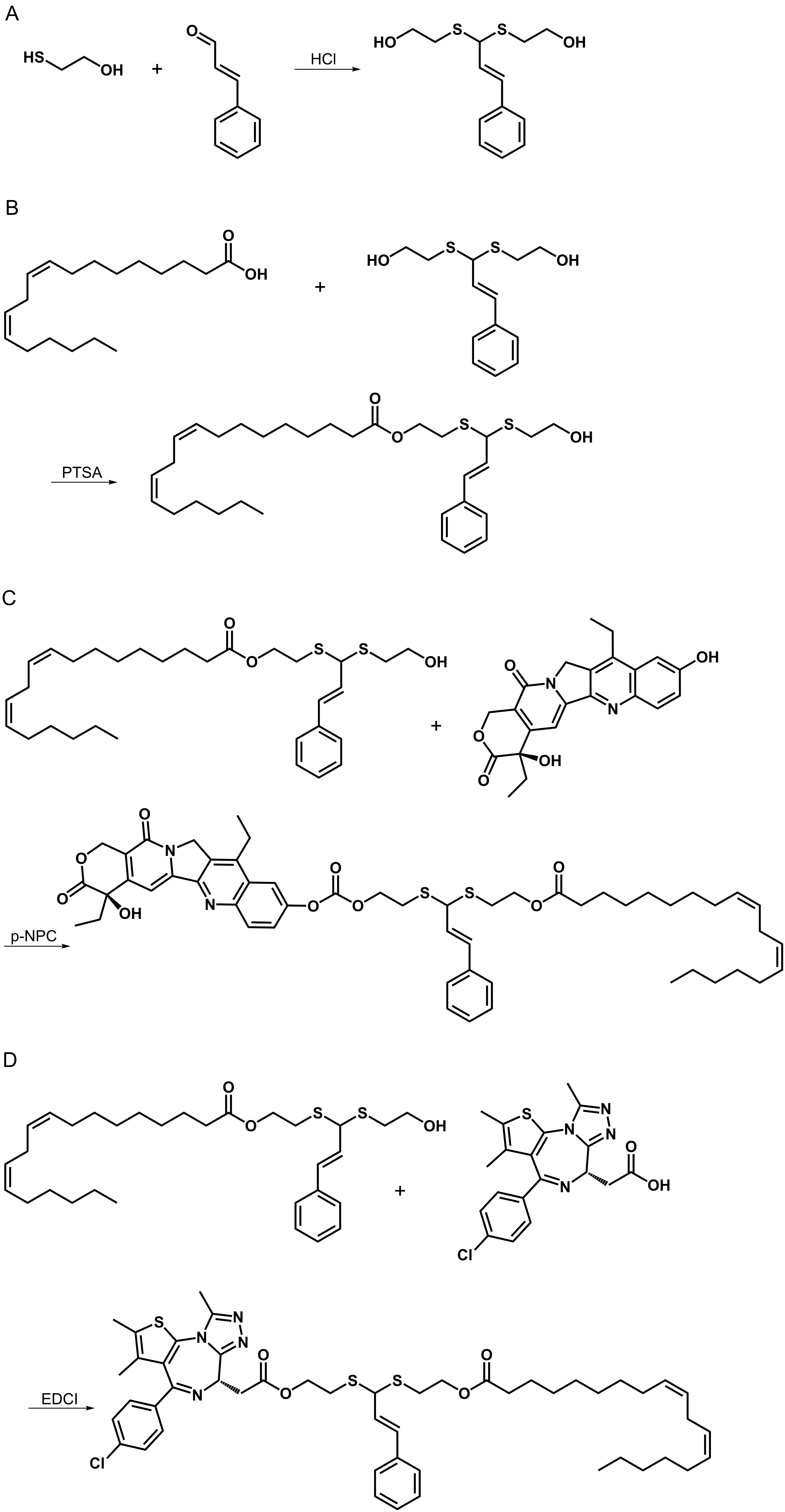

**Fig. S1. Chemical synthesis route.** A) Synthesis route of CT. HCl: hydrochloric acid. B) Synthesis route of LA-CT. PTSA: p-toluenesulfonic acid. C) Synthesis route of SN38-LA. P-NPC: p-nitrophenyl chloroformate. D) Synthesis route of JQ-1-LA. EDCI: 1-ethyl-3(3-dimethylpropylamine) carbodiimide.

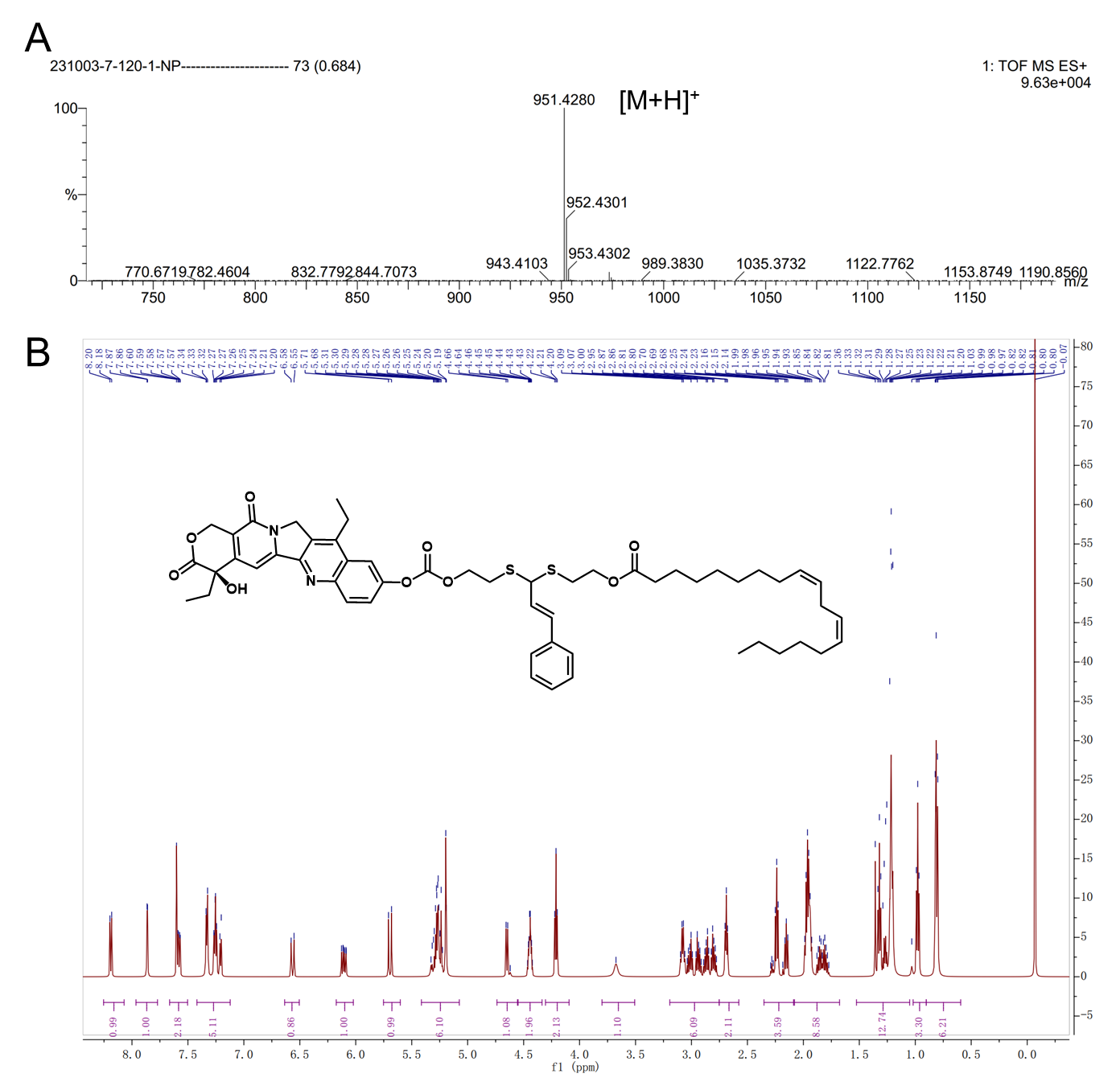

**Fig. S2. Structure conformation of SN38-LA.** A) MS. MS (ESI) m/z: Calcd for C_54_H_66_N_2_O_9_S_2_, 950.4210; found, 951.4280 [M+H]^+^. B) ^1^H NMR. ^1^H NMR (600 MHz, Chloroform-d) δ 8.19 (d, J = 9.1 Hz, 1H, Ar-H), 7.86 (d, J = 2.5 Hz, 1H, Ar-H), 7.66 – 7.50 (m, 2H, Ar-H), 7.42 – 7.12 (m, 5H, Ar-H, -N-C=CH-), 6.56 (d, J = 15.3 Hz, 1H, Ar-CH=CH-), 6.17 – 6.02 (m, 1H, Ar-CH=CH-), 5.41 – 5.07 (m, 6H, -CH=CH-, -O-CH_2_-), 4.65 (d, J = 8.6 Hz, 1H, -OH), 4.44 (td, J = 6.6, 3.3 Hz, 2H, -OCOO-CH_2_-), 4.21 (t, J = 6.8 Hz, 2H, -COO-CH_2_-), 3.80 – 3.51 (m, 1H, -CH-), 3.19 – 2.75 (m, 6H, N-CH_2_-, -CH_2_-CH_3_, -CH=CH-CH_2_-CH=CH-), 2.69 (t, J = 6.8 Hz, 4H, -S-CH_2_-), 2.20 (dt, J = 51.8, 7.6 Hz, 2H, -CO-CH_2_-), 2.08 – 1.68 (m, 8H, -CH=CH-CH_2_-, -CH_2_-CH_3_, -CO-CH_2_-CH_2_-), 1.53 – 1.05 (m, 14H, -CH_2_-), 0.98 (t, J = 7.4 Hz, 3H, -CH_3_), 0.90 – 0.59 (m, 6H, -CH_3_).

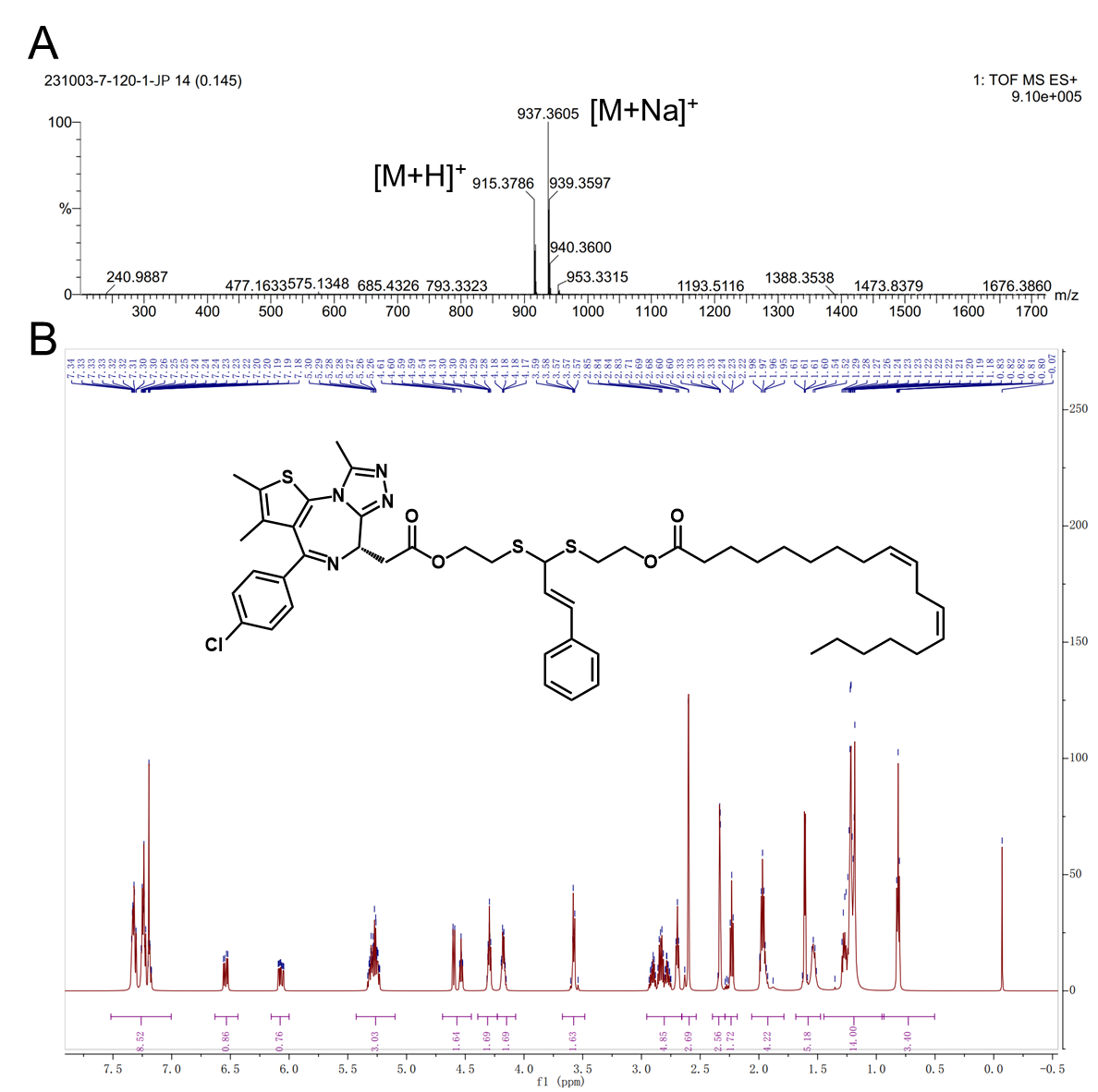

**Fig. S3. Structure conformation of JQ-1-LA.** A) MS. MS (ESI) m/z: Calcd for C_50_H_63_ClN_4_O_4_S_3_, 914.3700; found, 915.3786 [M+H]^+^ and 937.3605 [M+Na]^+^. B) ^1^H NMR. (600 MHz, Chloroform-d) δ 7.52 – 7.00 (m, 9H, Ar-H), 6.54 (dd, J = 15.6, 6.8 Hz, 1H, Ar-CH=CH-), 6.07 (ddd, J = 15.7, 8.7, 3.9 Hz, 1H, Ar-CH=CH-), 5.43 – 5.10 (m, 4H, -CH=CH-), 4.69 – 4.45 (m, 2H, JQ1-O-CH_2_-), 4.39 – 4.23 (m, 2H, LA-O-CH_2_-), 4.18 (tdd, J = 6.7, 4.2, 2.7 Hz, 2H, -CH-), 3.67 – 3.48 (m, 2H, -CO-CH_2_-(JQ1)), 2.95 – 2.66 (m, 6H, -S-CH_2_-, -CH=CH-CH_2_-CH=CH-), 2.60 (d, J = 1.4 Hz, 3H, -CH_3_), 2.40 – 2.29 (m, 3H, -CH_3_), 2.23 (t, J = 7.6 Hz, 2H, -CO-CH_2_-(LA)), 1.97 (p, J = 6.8 Hz, 4H, -CH=CH-CH_2_-), 1.69 – 1.48 (m, 5H, -CH_3_, -CO-CH_2_-CH_2_-), 1.45 – 0.94 (m, 14H, -CH_2_-), 0.95 – 0.50 (m, 3H, -CH_3_).

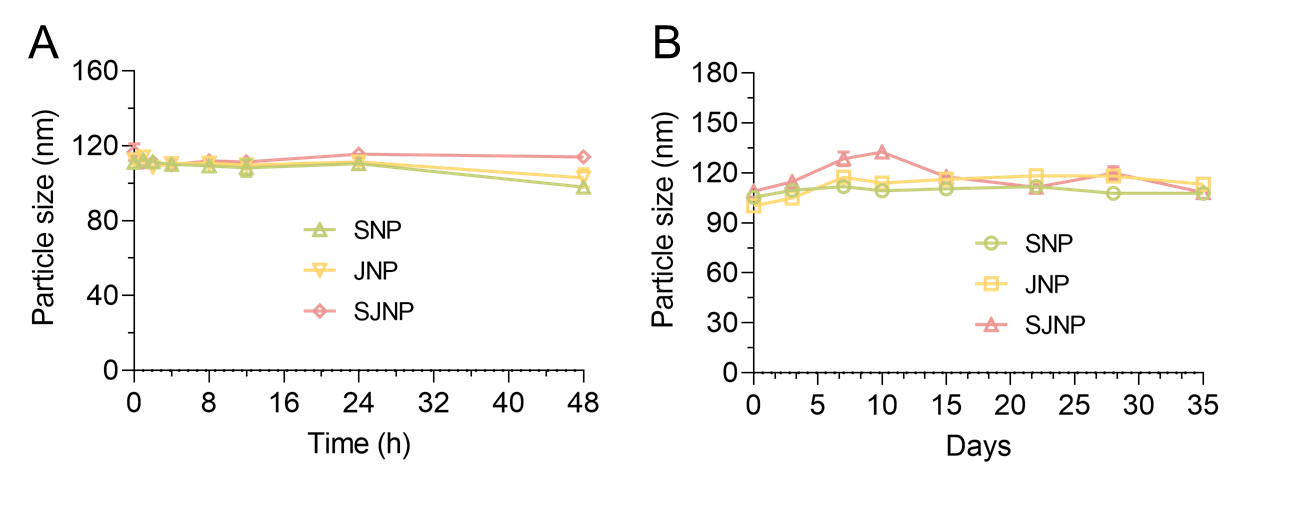

**Fig. S4. Colloidal stability.** A) Incubation of PANP in PBS (pH 7.4) supplemented with 10% FBS at 37℃ for 48 h. B) Storage of prodrug nanoassemblies in room temperature at 4℃ for 35 days. Data are portrayed as mean ± standard deviation (n = 3).

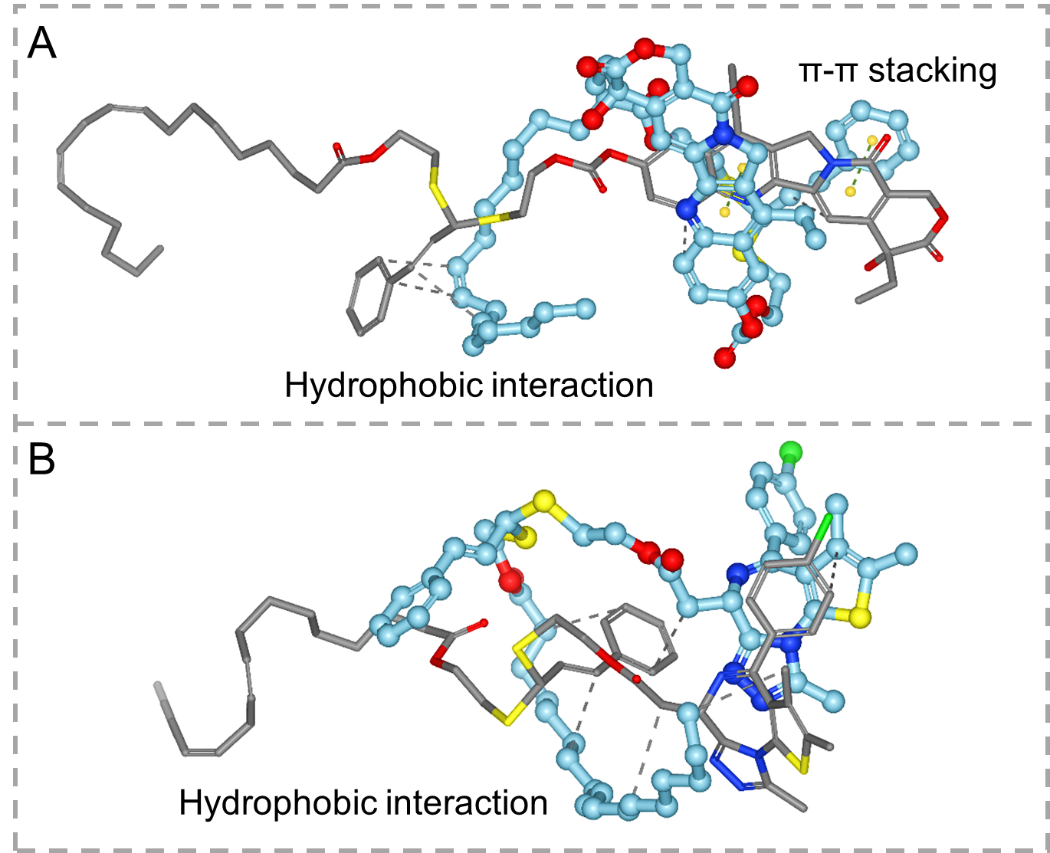

**Fig. S5. Molecular conformation.** A) Conformations of SNP during MD simulations. B) Conformations of JNP during MD simulations.

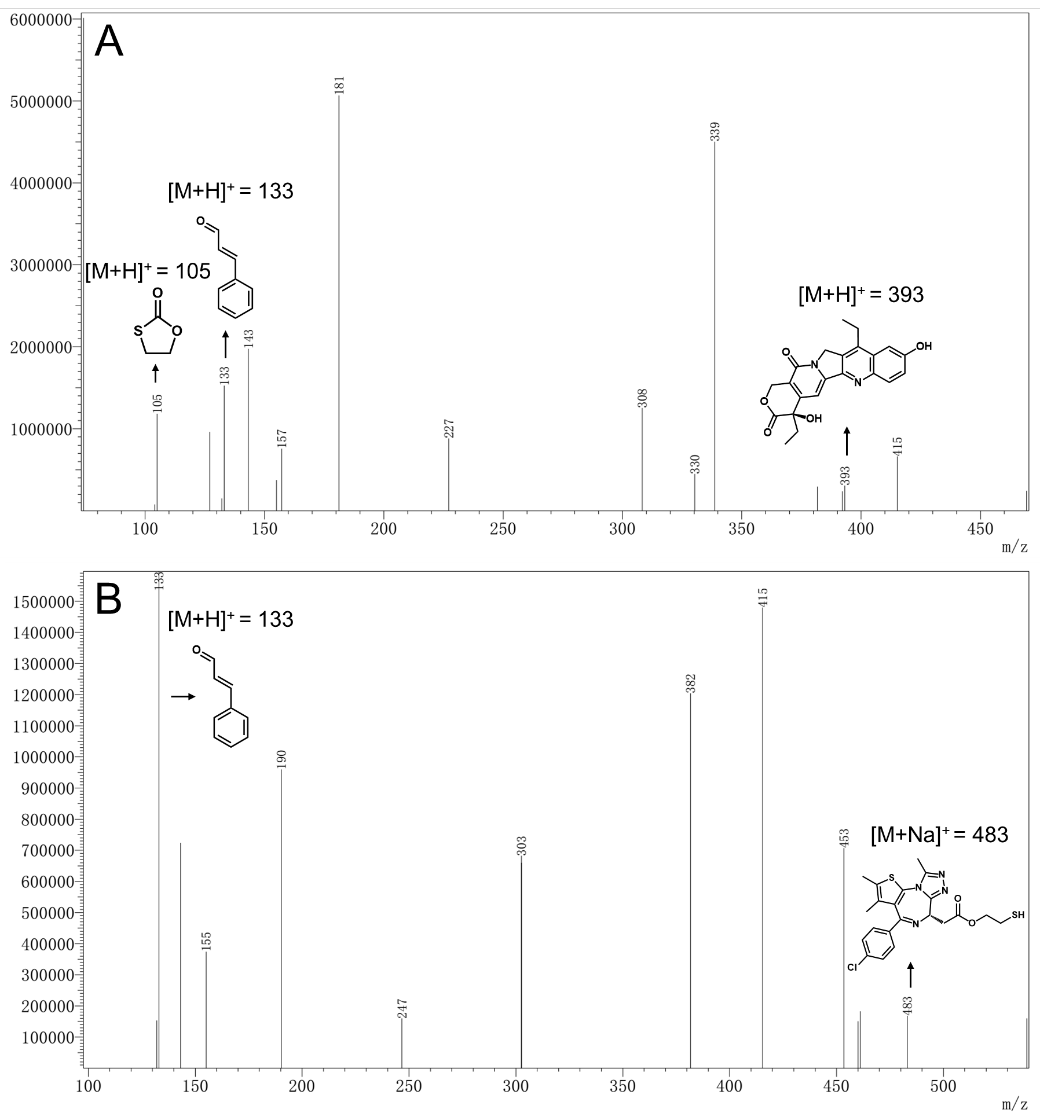

**Fig. S6. MS of release intermediates.** A) Release intermediates of SN38-LA including SN38, CA and 1,3-oxathiolan-2-one. B) Release intermediates of JQ-1-LA including JQ-1-SH and CA.

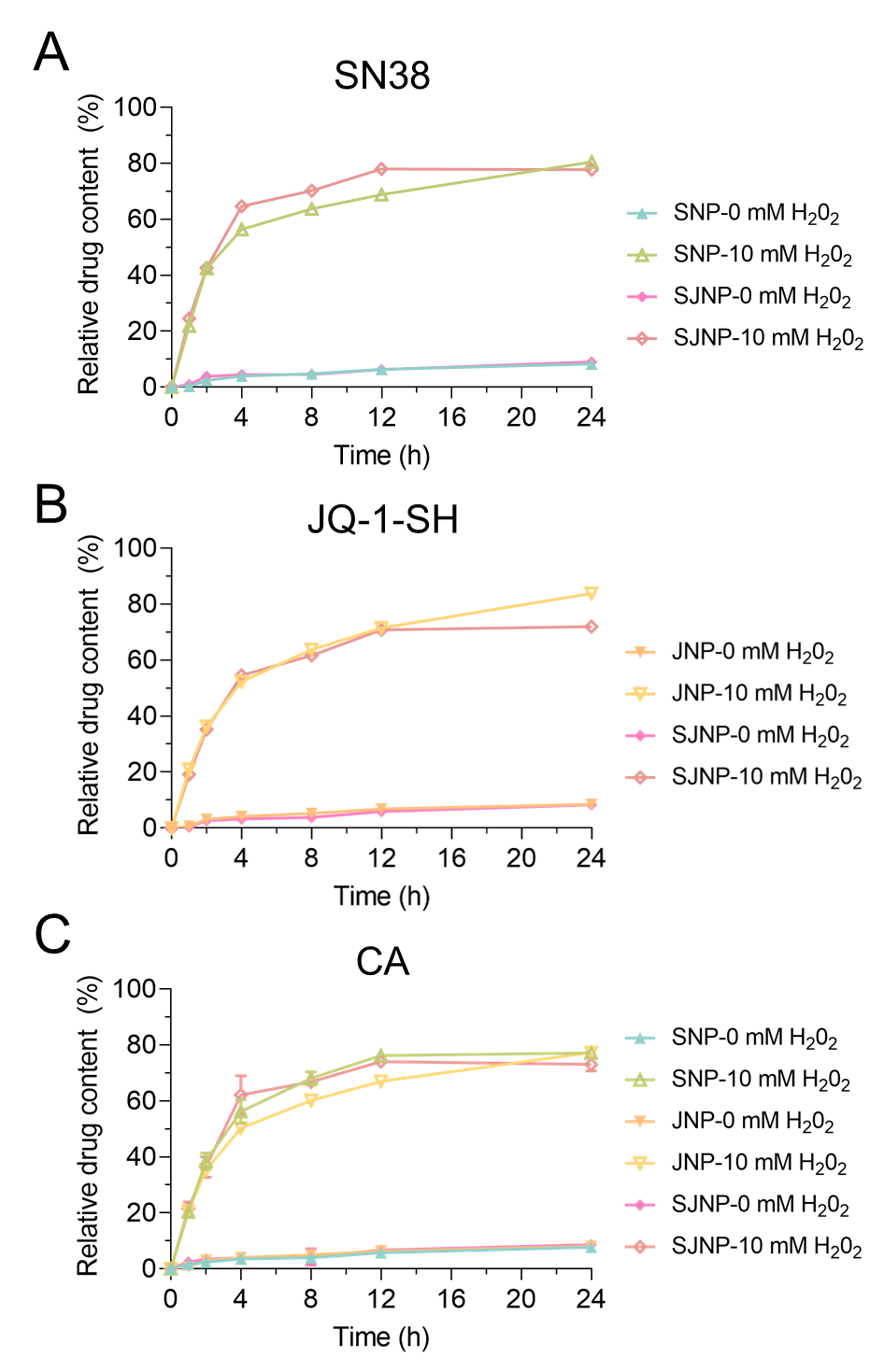

**Fig. S7. In vitro drug release of PANP with/without the presence of H_2_O_2_.** A) SN38 released from SNP and SJNP. B) JQ-1-SH released from JNP and SJNP. C) CA released from SNP, JNP and SJNP. Data are portrayed as mean ± standard deviation (n = 3).

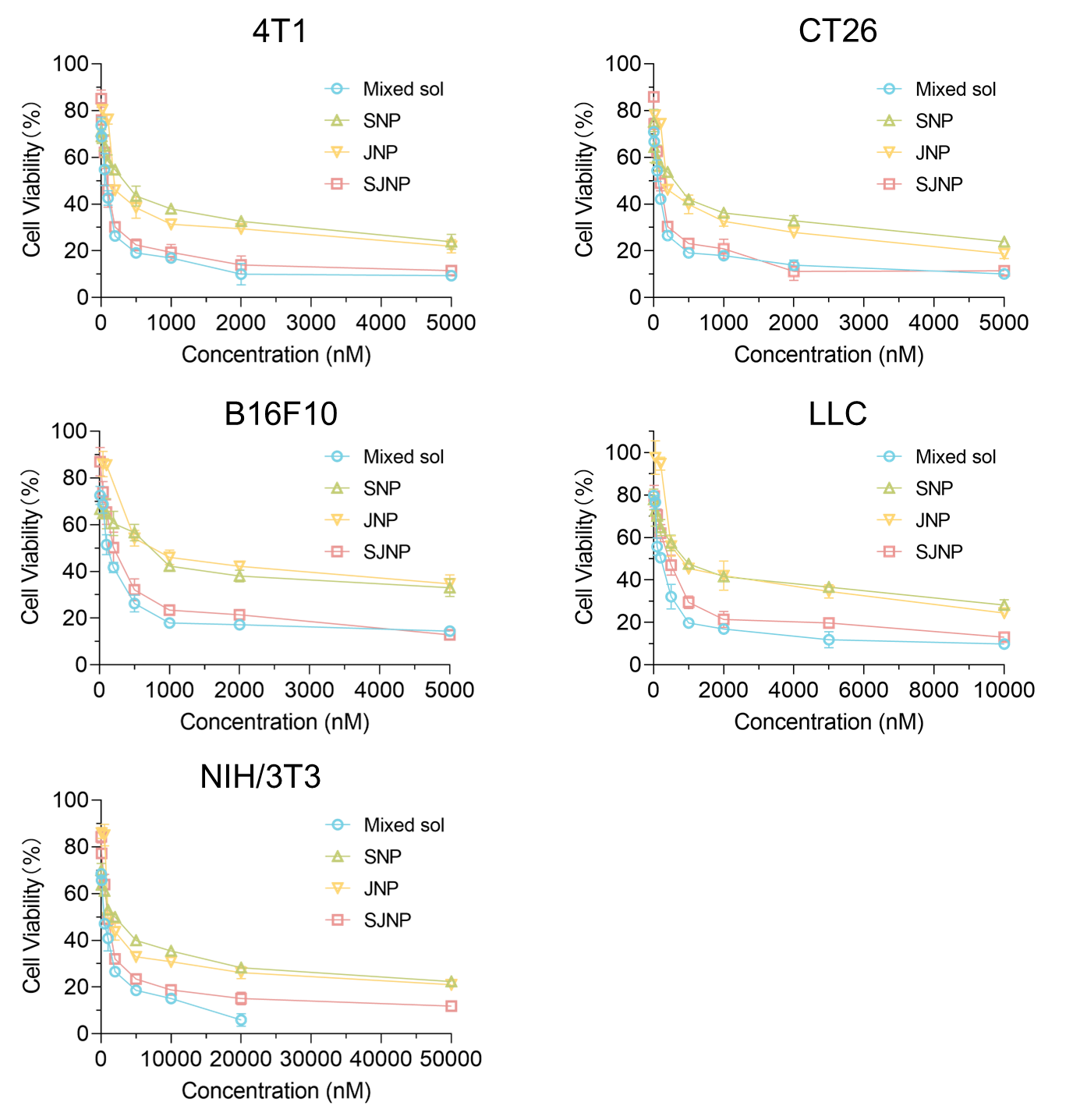

**Fig. S8. Cell viability of different cells post administration.** Data are portrayed as mean ± standard deviation (n = 3).

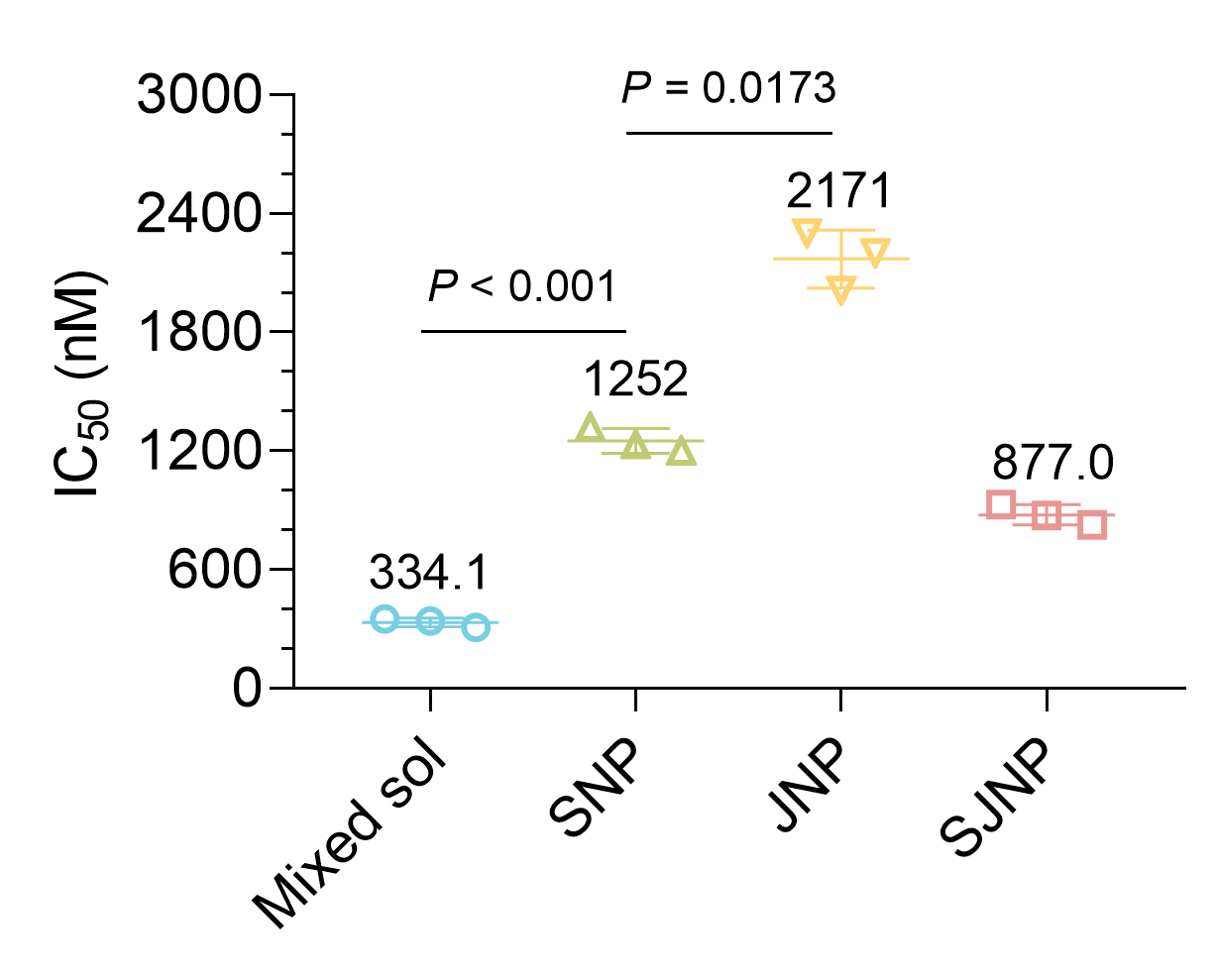

**Fig. S9. Cytotoxicity of PANP on NIH/3T3 cells.** Data are portrayed as mean ± standard deviation (n = 3). Statistical significance was assessed through ANOVA and deemed significant at P < 0.05.

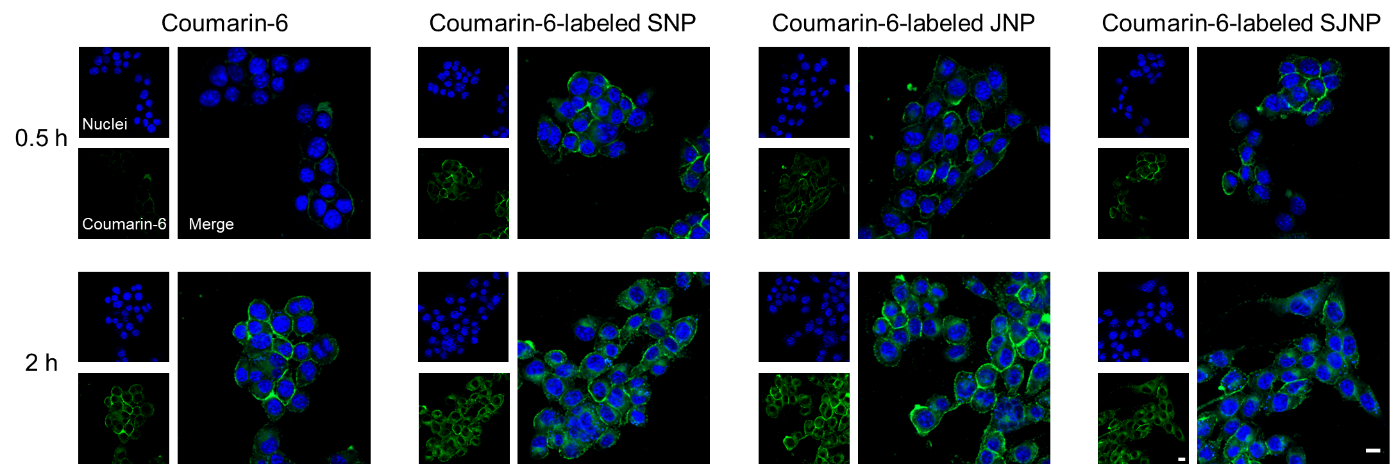

**Fig. S10. Cellular uptake of coumarin-6-labeled PANP on 4T1 cells.** Scale bar = 10 μm.

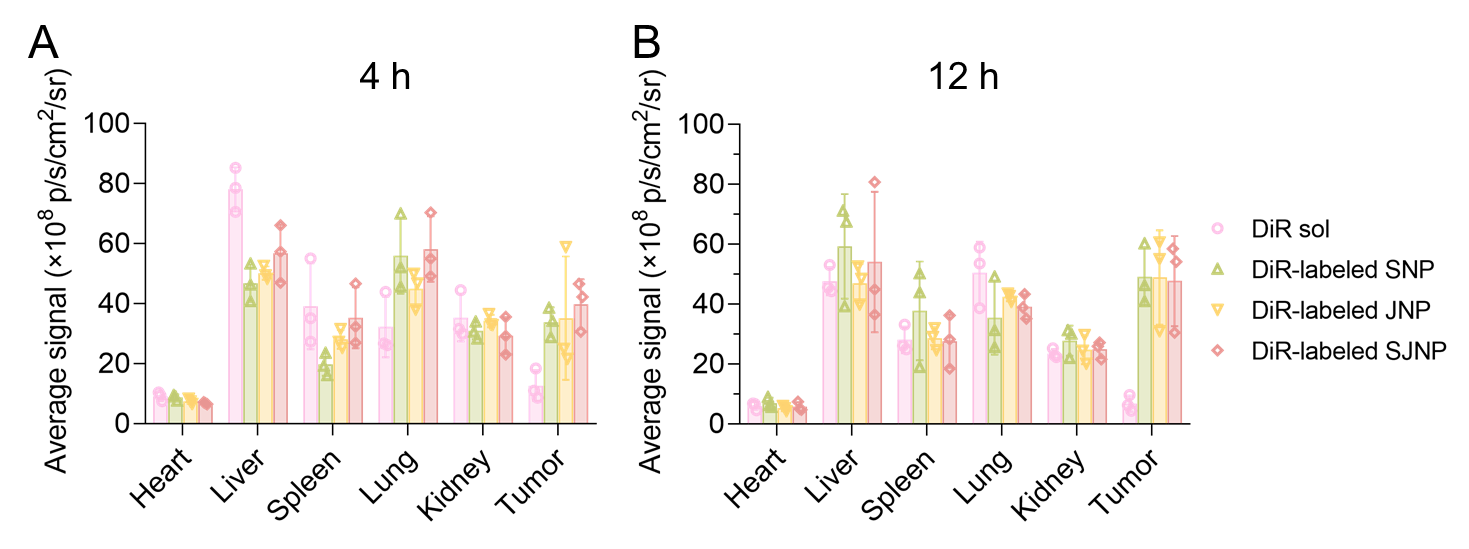

**Fig. S11. Quantitative results of biodistribution at A) 4 h and B) 12 h.** Data are portrayed as mean ± standard deviation (n = 3).

**Table S1. Characteristics of PANP.**

| Formulations | Size (nm) | PDI | ζ Potential (mV) | Drug loading (%) |
| --- | --- | --- | --- | --- |
| SNP | 83.45±0.139 | 0.143 | -39.9±0.723 | 32.9% |
| JNP | 100.5±0.289 | 0.118 | -36.9±1.81 | 34.9% |
| SJNP | 91.18±1.321 | 0.120 | -41.3±0.493 | 8.2% for SN38; 26.2% for JQ-1 |

Data are portrayed as mean ± standard deviation (n = 3).

**Table S2. Characteristics of co-assembled PANP.**

| Formulations^a)^ | Size (nm) | PDI |
| --- | --- | --- |
| 10:1 ratio | 90.90±2.335 | 0.113 |
| 5:1 ratio | 81.30±0.552 | 0.136 |
| 4:1 ratio | 86.10±0.778 | 0.077 |
| 3:1 ratio | 84.68±1.408 | 0.145 |
| 2:1 ratio | 93.88±3.257 | 0.137 |
| 1:1 ratio | 91.53±1.976 | 0.153 |
| 1:2 ratio | 93.79±0.910 | 0.227 |
| 1:3 ratio | 91.18±1.321 | 0.120 |
| 1:4 ratio | 94.11±1.480 | 0.179 |
| 1:5 ratio | 102.3±1.890 | 0.191 |
| 1:10 ratio | 104.5±1.286 | 0.177 |

a) Co-assembled PANP was prepared at various molar ratios (SN38-LA to JQ-1-LA). Data are portrayed as mean ± standard deviation (n = 3).

**Table S3. IC_50_ values of co-assembled PANP at different molar ratios on 4T1 cells.**

| Formulations^a)^ | IC_50_ (nM) |
| --- | --- |
| SN38 | 22.41 |
| JQ-1 | 7064 |
| SN38-LA | 139.6 |
| JQ-1-LA | 361.5 |
| 10:1 ratio | 254.2 |
| 5:1 ratio | 325.5 |
| 4:1 ratio | 210.6 |
| 3:1 ratio | 333.4 |
| 2:1 ratio | 263.3 |
| 1:1 ratio | 189.5 |
| 1:2 ratio | 191.8 |
| 1:3 ratio | 78.55 |
| 1:4 ratio | 254.3 |
| 1:5 ratio | 131.1 |
| 1:10 ratio | 227.9 |

a) Co-assembled PANP was prepared at various molar ratios (SN38-LA to JQ-1-LA).

**Table S4. SI of PANP on different tumor cells**

| Formulations | 4T1 | CT26 | B16F10 | LCC |
| --- | --- | --- | --- | --- |
| Mixed sol | 7.16 | 7.88 | 3.10 | 1.94 |
| SNP | 5.44 | 7.15 | 2.76 | 1.37 |
| JNP | 7.03 | 7.31 | 1.94 | 1.69 |
| SJNP | 11.14 | 10.63 | 4.06 | 2.39 |
